# A robotic system for automated genetic manipulation and analysis of *Caenorhabditis elegans*

**DOI:** 10.1101/2022.11.18.517134

**Authors:** Zihao Li, Anthony D. Fouad, Peter D. Bowlin, Yuying Fan, Siming He, Meng-Chuan Chang, Angelica Du, Christopher Teng, Alexander Kassouni, Hongfei Ji, David M. Raizen, Christopher Fang-Yen

## Abstract

The nematode *Caenorhabditis elegans* is one of the most widely studied organisms in biology due to its small size, rapid life cycle, and manipulable genetics. Research with *C. elegans* depends on labor-intensive and time-consuming manual procedures, imposing a major bottleneck for many studies, especially those involving large numbers of animals. Here we describe the first general-purpose tool, WormPicker, a robotic system capable of performing complex genetic manipulations and other tasks by imaging, phenotyping, and transferring *C. elegans* on standard agar media. Our system uses a motorized stage to move an imaging system and a robotic arm over an array of plates. Machine vision tools identify animals and assay developmental stage, morphology, sex, expression of fluorescent reporters, and other phenotypes. Based on the results of these assays the robotic arm selectively transfers individual animals using an electrically self-sterilized wire loop, with the aid of machine vision and electrical capacitance sensing. Automated *C. elegans* manipulation shows reliability and throughput comparable to standard manual methods. We developed software to enable the system to autonomously carry out complex protocols. To validate the effectiveness and versatility of our methods we used the system to perform a collection of common *C. elegans* procedures, including genetic crossing, genetic mapping, and genomic integration of a transgene. Our robotic system will accelerate *C. elegans* research and opens possibilities for performing genetic and pharmacological screens that would be impractical using manual methods.

**Significance Statement:** The nematode *Caenorhabditis elegans* is a powerful genetic model organism in life sciences due to its compact anatomy, short life cycle, and optical transparency. Current methods for worm genetics rely on laborious, time-consuming, and error-prone manual work. Here, we describe the first general-purpose automated tool capable of genetically manipulating *C. elegans*. Our robotic system will accelerate a broad variety of *C. elegans* research and opens possibilities for performing genetic and pharmacological screens that would be impractical using manual methods.

## Introduction

Classical genetics, which investigates the heritability of traits across generations, usually requires manipulating reproductive behaviors of organisms and inferring their genetic properties by assaying their traits. The microscopic nematode *C. elegans* is one of the most widely used genetic models in life sciences due to its easy maintenance, optical transparency, and rapid life cycle (1, 2). Studies in *C. elegans* have pioneered major fundamental advances in biology, including those in programmed cell death (3), aging (4), RNA interference (5), and axon guidance (6). Work with worms has pioneered important techniques in modern biology including genome sequencing (7), cell lineage tracing (3), gene editing (8), electron microscopic reconstruction of neural connectivity (9), demonstration of green fluorescent protein (10), and optogenetic manipulation of neural activity (11).

Genetic manipulations in *C. elegans* are performed by manual procedures, which involve identification of animals under a microscope and transfer of worms or embryos from one agar plate to another using a wire pick (12). While manual procedures are reliable and technically simple, they have important limitations.

Manual methods are labor intensive since they require animals to be manipulated individually. This presents challenges to experiments that require thousands of groups to be managed, for example conducting genetic screens (13), working with collections of wild isolates (14), or dealing with mutagenized strains of the Million Mutation Project (15).

During design of lab experiments, the use of manual procedures creates practical limits to the number of conditions and the number of replicates for each condition, weakening statistical power. Standard population sizes used for *C. elegans* lifespan experiments have been shown to be underpowered for moderate differences in lifespan between groups (16).

Finally, manual approaches require training and are prone to errors. This reliance on investigator-learned skills imposes a barrier to entry for scientists without *C. elegans* experience wishing to use this system.

To address the limitations of manual methods, automated organism manipulations have been reported for bacteria and single cells in the field of synthetic biology (17) and for adult *Drosophila* (18). For work with *C. elegans*, automated imaging systems have been developed for behavioral analysis (19), lifespan or healthspan measurement (20–22), and drug screening (23). Microfluidic (24–28) and flow-cell devices (29, 30) have been demonstrated for high-throughput animal imaging and sorting. However, to our knowledge no methods for automated genetic manipulations in *C. elegans* have been reported.

Here we present WormPicker, a general-purpose robotic system allowing automated phenotyping and genetic manipulation of *C. elegans* on agar substrates, using techniques resembling manual methods. Our device contains a 3D motorized stage carrying a robotic arm and an optical system. The robotic arm manipulates animals using a thin, electrically sterilized platinum wire loop. Analogous to manual methods, the robotic arm picks worms by performing spatially and temporally controlled motions above and on the agar surface, using food bacteria to encourage the worm to adhere to the loop. Contact between the platinum wire loop and the agar surface is perceived by a capacitive touch sensing circuit, providing feedback in conjunction with the imaging system for fine adjustment of the pick trajectory relative to the animal.

The robot’s optical system is capable of monitoring animals over an entire plate at low magnification (6 cm diameter circular field of view (FOV)) while simultaneously imaging individual animals at high magnification (1.88 mm x 1.57 mm FOV) to obtain more detailed morphological and/or fluorescence information. Using machine vision methods, worms can be recognized and tracked over the plates in low magnification and undergo detailed phenotyping in high magnification across different attributes, including developmental stage, morphology, sex, and fluorescence expression. We developed system control software through which the user can specify multi-step genetic procedures to be performed.

Using these automated tools, we successfully carried out three genetic procedures commonly performed in *C. elegans* research. First, we generated a genetic cross between transgenic and mutant animals using a classic genetic hybridization scheme. Second, we performed genetic mapping of a genome-integrated fluorescent transgene. Finally, we integrated an extrachromosomal transgenic array to the genome, creating a stable transgenic line. Successful completion of these complex genetic procedures demonstrates WormPicker’s effectiveness and versatility as a broadly useful tool for *C. elegans* genetics.

## Results

### Overview of WormPicker system design

Our system contains a robotic picking arm, optical imaging system, lid manipulators, and other elements mounted on a 3-dimensional motorized stage to work with an array of up to 144 agar plates (Fig. 1*A*, *B*, and *SI Appendix*, Fig. S1).

**Fig. 1.**
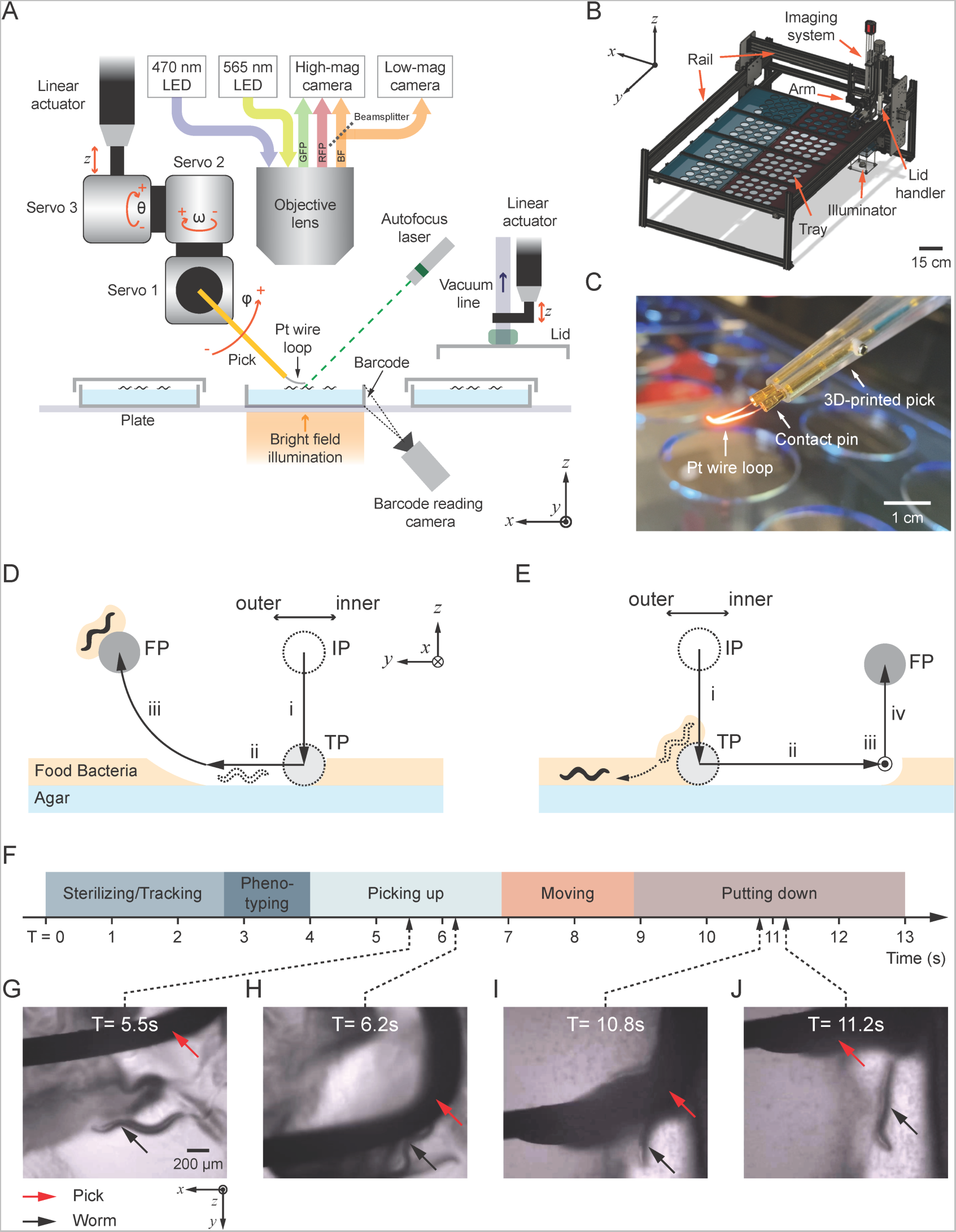
WormPicker system configuration. *(A)* WormPicker geometry. *C. elegans* is manipulated by a wire pick motorized by servo motors and a linear actuator. Animals are imaged by an optical imaging stream. Lids are manipulated by a vacuum grabbing system. The imaging and manipulation assemblies are mounted on a 3D motorized stage. *(B)* WormPicker system overview. *(C)* Photograph of the 3D-printed pick, captured when the wire loop being sterilized by an electric current. *(D* and *E)* Schematic of pick motion trajectories for *C. elegans* picked up from and put down to agar substrate. IP: initial position; TP: agar-touching position; FP: final position. *(F)* Timeline for stages throughout the automated transfer. *(G-J)* Key frames captured by the high-magnification camera during the Picking up *(G* and *H)* and Putting down stage *(I* and *J)*.

The array of plates is housed on a platform (Fig. 1*B*), containing 8 trays that can be prepared separately and then slid into the automated system (*SI Appendix*, Fig. S2*A* and *B*). Each plate on the tray can be labeled with an adhesive barcode and text (Fig. 1*A* and *SI Appendix*, Fig. S2*C* and *D*). We developed software to track plates during experiments using their barcode identifiers (*SI Appendix*, Fig. S2*G-I*).

**Fig. 2.**
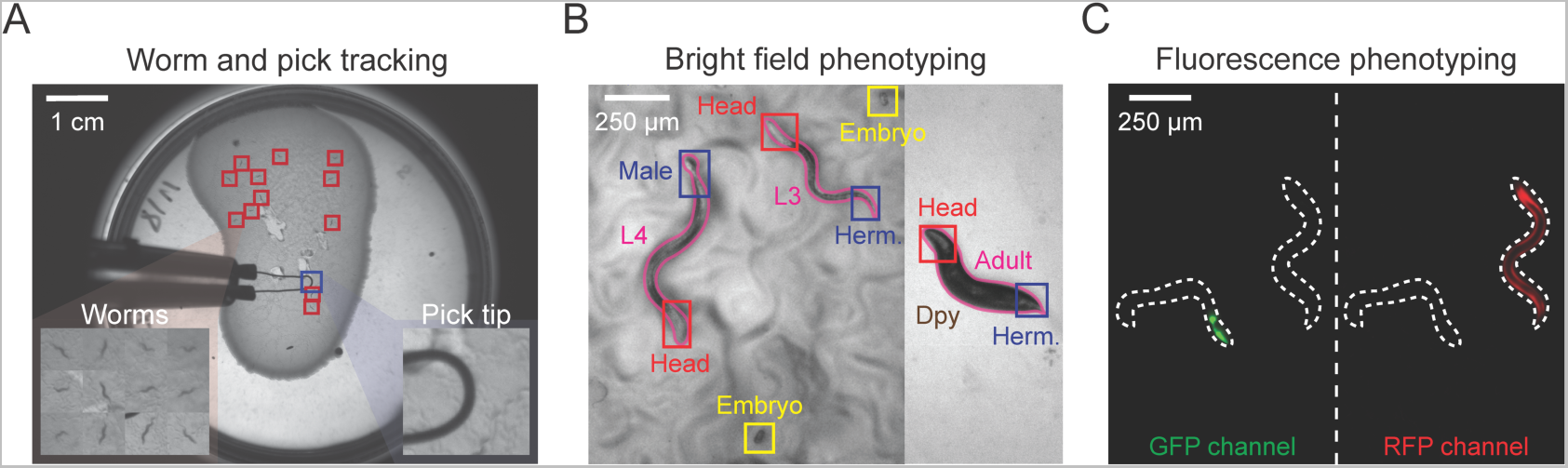
Overview of machine vision approaches. Machine vision algorithms were developed for analyzing frames acquired from the *(A)* low and *(B* and *C)* high-magnification imaging streams. The low-magnification frames *(A)* are analyzed for tracking worms and the pick in real time. The high-magnification bright field images *(B)* are processed by a set of Mask-Regional CNNs (Mask-RCNNs) for close inspection of individual animals, including contour segmentations (pink solid line) and assays of developmental stage (including embryo), sex, and morphology. The high-magnification fluorescent images *(C)* are subject to intensity analysis for identifying fluorescent markers in GFP and RFP channels. White dashed line: animal contours segmented in the bright field.

The imaging system is designed for bright field and fluorescence imaging through low (0.035X) and high-magnification (10X) optical pathways. The low-magnification stream has a 6 cm diameter circular FOV, capable of imaging one 6 cm diameter plate; the high-magnification stream has a 1.88 mm x 1.57 mm FOV, capable of imaging individual animals in detail. For bright field imaging, light is collected by an objective lens and divided into low and high-magnification imaging streams by a beamsplitter (Fig. 1*A* and *SI Appendix*, Fig. S3*A*). For fluorescence imaging, collimated excitation LEDs (center wavelengths 470 nm and 565 nm) and a fluorescence optics (*SI Appendix*, Fig. S3*B* and *C*) enable imaging in green (503 – 541 nm wavelength) and red (606 – 661 nm wavelength) channels (*SI Appendix*, Movie S1) via the high-magnification pathway.

**Fig. 3.**
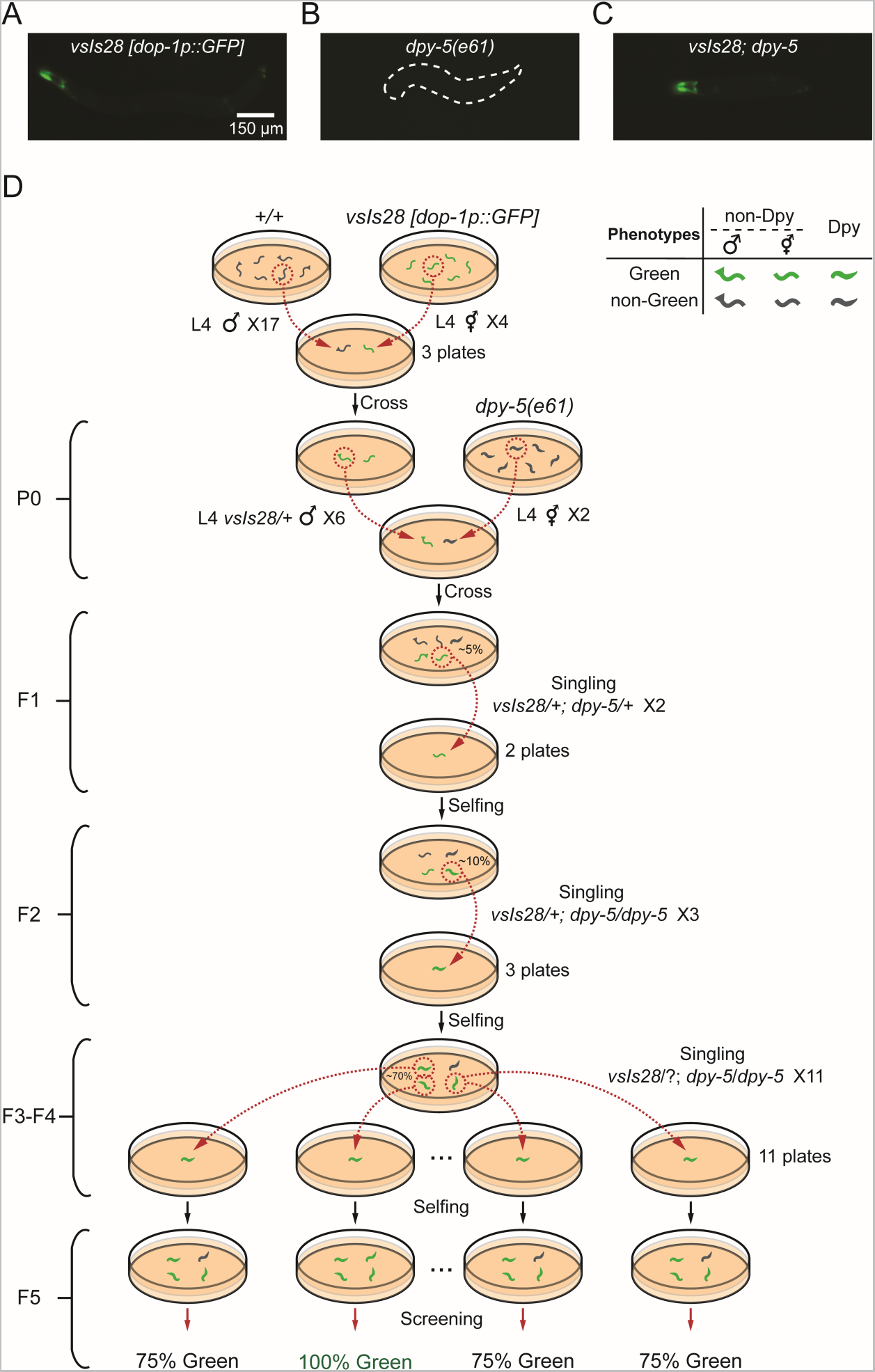
Automated genetic cross in *C. elegans*. WormPicker generated a genetic cross between animals carrying an integrated fluorescent transgene *vsIs28 [dop-1p::GFP]* and *dpy-5(e61)* mutants. *(A-C)* GFP expression in the strain carrying *(A) vsIs28*, *(B) dpy-5*, and their hybridization *(C) vsIs28*; *dpy-5*. *(D)* Schematic of the genetic cross.

To quickly bring the surface of a plate into focus, we use a laser-based autofocusing system (Fig. 1*A* and *SI Appendix*, S3*D*). A 532 nm laser pointer with an integrated cylindrical lens generates a line projected onto the agar surface from an oblique angle, providing feedback for the motorized stage to move the imaging system to the correct position.

Plate lids are manipulated by a vacuum gripping system (Fig. 1*A* and *SI Appendix*, Fig. S3*E*). Two lid handlers are mounted to either side of the main motorized gantry for manipulating lids for a source plate and a destination plate separately (only one visible in Fig. 1*A*). Each vacuum lid gripper is raised and lowered by a motorized linear actuator (*SI Appendix*, Movie S2).

The robotic picking arm (Fig. 1*A* and *SI Appendix*, Fig. S3*F*) contains a motorized linear actuator for fine adjustment of the pick’s height and three servo motors to provide rotational degrees of freedom. The 3D-printed pick contains a platinum wire loop attached to its end (Fig. 1*C*), for manipulating *C. elegans* in a manner analogous to manual methods (12).

Conventional pick sterilization prior to manipulating *C. elegans* on agar substrates requires an open flame, which poses safety risks in an automated system. We adopted an electric sterilization approach (Fig. 1*C*) in which a current is passed through the wire loop to sterilize it via resistive heating (*SI Appendix*, Movie S3).

Picking up *C. elegans* requires very fine control of the pick to avoid damage to the worm or the agar surface. While the horizontal position of the pick can be monitored by the imaging system, its height relative to the agar surface is more difficult to determine. To address this problem, we developed a capacitive touch sensing circuit (*SI Appendix*, Fig. S3*G*) that monitors contact between the platinum wire and agar surface and provides feedback for fine tuning the pick’s movements.

To pick up a worm (Fig. 1*D*), the wire pick is positioned above the agar (IP), with a horizontal offset to the target worm. The linear actuator lowers the pick until contacting the agar surface (TP) as perceived by the capacitive touch sensor (phase i). Next, Servo 2 horizontally swipes the pick on the agar surface (phase ii). Servos 1 and 3 act simultaneously to perform a curved motion (phase iii) for picking up the target animal using the outer side of the wire loop. To release the worm from the pick to the substrate (Fig. 1*E*), the pick is positioned above the substrate (IP); the linear actuator vertically lowers (phase i) the pick until it touches the agar (TP); the Servo 2 horizontally swipes (phase ii) the pick on the agar surface horizontally, to release the worm from the pick. *SI Appendix*, Movie S4 depicts these operations as observed by the WormPicker imaging system; *SI Appendix*, Movie S5 depicts these actions as observed by an external camera.

### Machine vision enables automated identification and phenotyping

We developed machine vision analysis software for the low and high-magnification streams. We analyze the low-magnification images (Fig. 2*A*) using a combination of a convolutional neural network (CNN) and motion detection for tracking animals and the pick in real time. High-magnification bright field images (Fig. 2*B*) are analyzed by a set of Mask-Regional CNNs (Mask-RCNNs) (31) capable of performing pixel-wise object segmentations. Animals’ contour geometries are analyzed for developmental stage and morphology. In addition, we trained separate networks for sex determination and embryo detection. Inferences from multiple Mask-RCNNs are integrated for assaying phenotypes for individual animals over different attributes.

For high-magnification fluorescence imaging (Fig. 2*C*), we apply intensity analysis, through which valid fluorescent signals are extracted from the background. To analyze fluorescence expression, the fluorescence images are correlated with the animal contours segmented using the bright field images (Fig. 2*C*).

Details of our machine vision approaches are given in *SI Appendix*, Extended Methods and Fig. S5.

### WormPicker reliably picks *C. elegans* of various stages and phenotypes

First, we asked whether the automated picking causes damage to *C. elegans*. We used our system to pick animals of all stages, ranging from L1 larvar to day 5 adults, and an array of mutants, including *lin-15* (multivulva), *rol-6* (roller), *unc-13* (uncoordinated and paralyzed), and *lpr-1* (fragile cuticles) (32). We measured the number alive 24 hours after automated picking. As control, we repeated the procedure using the standard manual methods (12). We observed that animals manipulated by WormPicker showed viability comparable to that of standard methods (*SI Appendix*, Table S1).

We asked how effectively the system picks up and puts down animals. We manually verified the success of individual pick-up and put-down attempts through the live image stream. We observed success rates about 90% for picking up and putting down different types of animals (*SI Appendix*, Table S2). When working with unseeded plates we precoated the wire loop with bacteria and observed similar success rates as for seeded plates.

These experiments show that WormPicker is safe and effective for transferring many different types of *C. elegans*, including young, aged animals, and various mutants.

### Automated throughput is comparable to that of experienced researchers

To compare the rate of automated and manual picking, we evaluated how quickly the robot and human researchers could perform a fluorescent animal sorting task. We used a strain in which some but not all worms carry a *myo-2::GFP*-labeled fluorescent extrachromosomal array (YX256); the task was to sort these worms into two plates containing fluorescent and non-fluorescent animals.

The robotic system sorted the animals (mixed stages) with a throughput of 3.21 ± 0.66 (mean ± SD) animals per minute (APM).

We recruited a group of *C. elegans* researchers (N = 21) to perform the same task using standard methods (12). The mean and median years of their *C. elegans* experience were 7.61 and 5 years, respectively. We tasked each volunteer to sort approximately 20 animals (of mixed stages) under a fluorescence stereoscope. Both WormPicker and the researchers picked individual worms and sterilized the pick between transfers. The measured throughputs are plotted versus years of experience in *SI Appendix*, Fig. S6. The manual picking throughput was 3.56 ± 1.67 (mean ± SD) APM.

These results show that the throughput of the robotic system for this fluorescent animal sorting task is comparable to that of experienced human researchers.

### The automated system maintains an aseptic environment

As in manual work, it is important to minimize contamination of media in our automated system. We designed the WormPicker to maintain an aseptic environment. We built the system inside a panel enclosure to prevent airborne contaminants from entering. We sanitized the active components, including the robotic arm, microscope, and plate trays using 70% ethanol before experiments. During experiments, plates had lids on for most of the time, except briefly during picking operations. For the experiments reported in *SI Appendix*, Tables S1 and S2, we did not observe any plate contaminated 10 days after being manipulated by WormPicker (N = 101 plates). In comparison, 3.4% of the plates were contaminated after manual picking in the same room (N = 119 plates). These results show that our protocols were sufficient for maintaining an aseptic environment for the experiments.

### Scripting toolsets enable complex genetic manipulations

In order for WormPicker to be useful for practical laboratory work, the basic elements of identifying and transferring worms need to be combined to form complex genetics procedures. To that end, we developed system control software, WormPickerControl (*SI Appendix*, Fig. S7*A*). An Application Programming Interface (API) enables the user to specify *C. elegans* procedures to be carried out by the automated system. We developed a library of source scripts, each responsible for a specific task ranging from simple to complex. The automated system catalogs a set of plates based on their barcode s (Fig. 1*A* and *SI Appendix*, Fig. S2) and stores the information in a database.

Using WormPickerControl, we developed and tested a collection of genetic procedures commonly undertaken in *C. elegans* labs (Figs. 3-5). These are not intended to be a complete set of genetic protocols, but rather to provide a useful framework that can be modified and adapted to other experiments as needed.

**Fig. 5.**
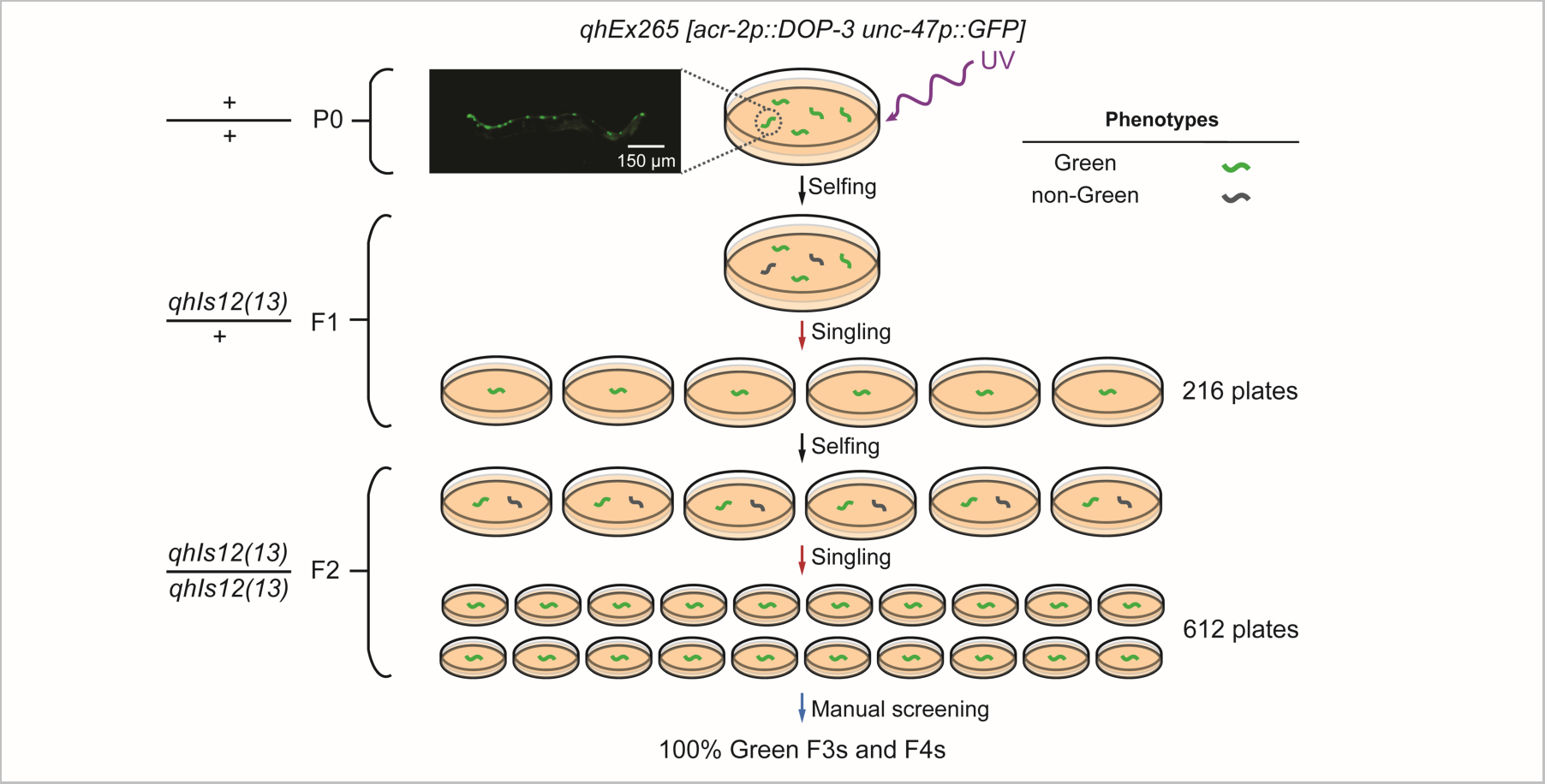
Automated genomic integration of a green fluorescence-labeled extrachromosomal array *qhEx265 [acr-2p::DOP-3 unc-47p::GFP]*. The image shows the GFP expression of the array. *qhIs12* and *qhIs13*: names of the potential integrated transgenic alleles.

### Automated genetic cross

The genetic cross, by which two or more mutations or transgenes are combined, is performed by placing males together with hermaphrodites on the same agar plate (33). While monitoring the plates, researchers pick cross-progeny with desired phenotypes. Most genetic crosses require manipulating animals over multiple generations.

We automated a genetic cross between two *C. elegans* strains: *dop-1p::GFP* fluorescent transgenic (LX831) and a *dpy-5(e61)* mutant (CB61) (Fig. 3). The LX831 strain contains an integrated transgene *vsIs28 [dop-1p::GFP]* and has a Green phenotype, i.e., GFP is expressed in several cell types including some neurons and muscles (Fig. 3*A*). The *dpy-5* mutant has a Dumpy (Dpy) phenotype, characterized by a morphology that is shorter and stouter than wild-type animals (Fig. 3*B*). The *vsIs28* transgene is dominant (i.e., both heterozygotes and homozygotes show green fluorescence) whereas the *dpy-5* mutation is recessive (i.e., only the homozygotes show the Dpy phenotype). The resulting hybridized strain displays both phenotypes of two parental strains, i.e., it is both Green and Dpy (Fig. 3*C*).

Fig. 3*D* depicts a schematic of the genetic cross. WormPicker picked 17 L4 males from a wild-type (N2) population that consisted of a mixture of males and hermaphrodites. It also picked 4 L4 hermaphrodites from the LX831 strain to the same plates. Green males were visible in the progeny, indicating that the mating was successful. Next, six Green L4 *vsIs28* heterozygous males and two *dpy-5* homozygous L4 hermaphrodites were picked onto a plate for mating. We consider this mating the parental (P0) generation.

Filial generation 1 (F1) progeny included animals that were Green and not Dpy. These were the desired worms heterozygous for both *vsIs28* and *dpy-5* and were therefore transferred by WormPicker to fresh plates. The automated system screened over the plate and singled two F1 hermaphrodites with the wanted phenotypes.

Four phenotypes were observed in the F2 generation, including Dpy-Green, Dpy-non-Green, non-Dpy-Green, and non-Dpy-non-Green. The frequency of desired Dpy-Green phenotype was ∼10%. WormPicker inspected the F2 populations and singled 3 Dpy-Green animals.

F3 populations descending from each of the three F2s were 100% Dpy but only 75% Green indicating that they were homozygous for *dpy-5* but heterozygous for *vsIs28*. WormPicker then singled 11 Dpy-Green animals.

The automated system screened for percentage of Green F5s descending from each of eleven F4s and identified one line displaying homozygous fluorescent, i.e., 100% of F5s were Green.

The results were verified by an experienced *C. elegans* researcher by noting that the resulting strain was positive for both Green and Dpy phenotypes, and that these phenotypes bred true in subsequent generations.

### Automated genetic mapping of a transgene

The identification of the genotype causing a particular phenotype usually requires genetic linkage analysis. The first step in such analysis is the identification of the chromosomes harboring the genetic change.

*C. elegans* has six chromosomes, of which five are autosomes and one is the X chromosome. Hermaphrodites are diploid for all of six chromosomes, while males are diploid for five autosomes and haploid for the X chromosome.

Genetic mapping in *C. elegans* can be performed by setting up genetic crosses between the strain of interest and a set of marker strains, and measuring the segregation pattern between the marker phenotypes and the phenotype of interest.

We used WormPicker to perform an automated genetic mapping of an integrated red fluorescent transgene *vsIs33 [dop-3p::RFP]* (LX811) (Fig. 4). This strain has a Red phenotype, i.e., RFP is expressed in cells expressing DOP-3 dopamine receptors (Fig. 4*A*).

**Fig. 4.**
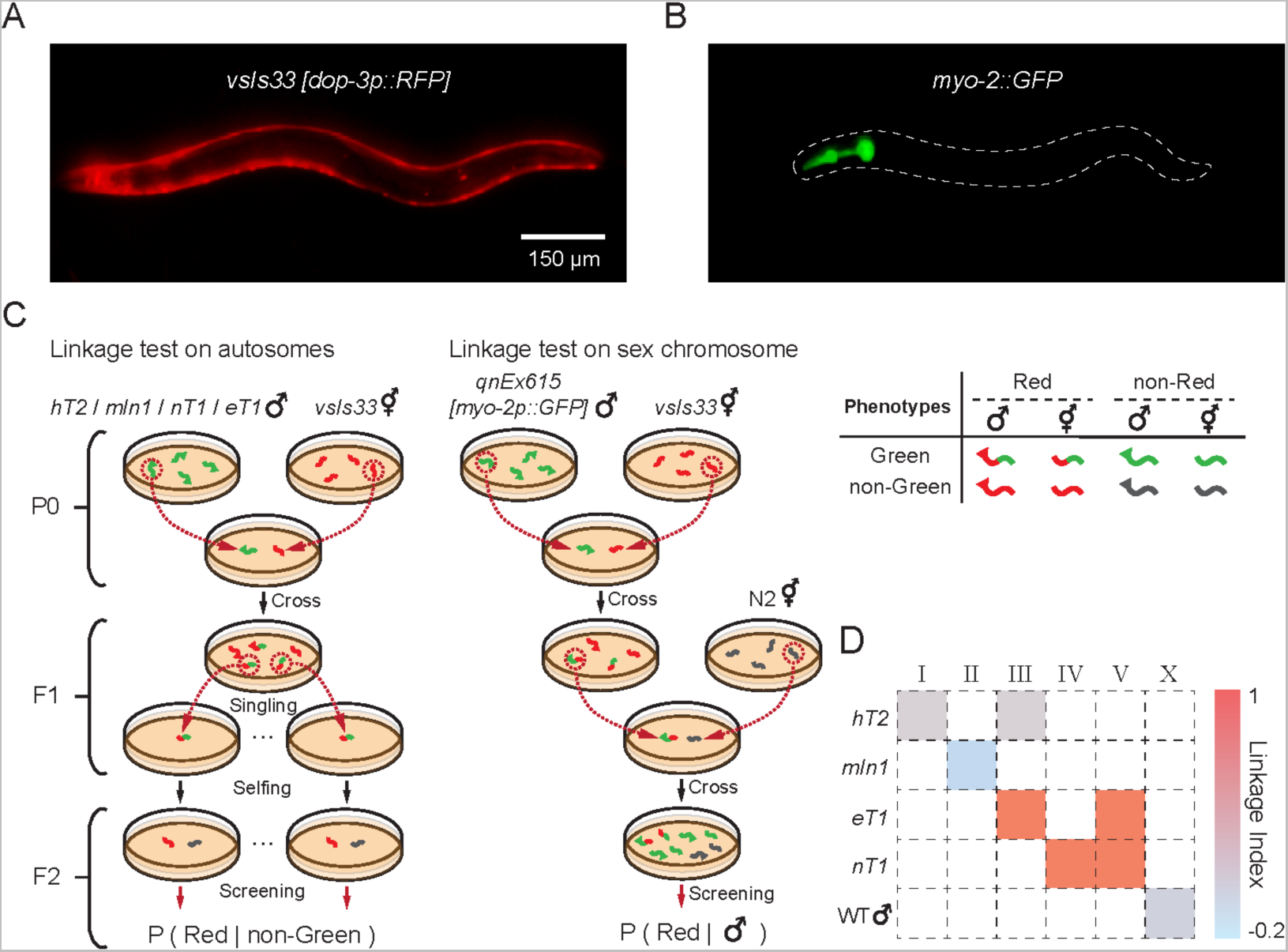
Automated genetic mapping of a red fluorescent transgene in *C. elegans*. *(A)* RFP expression in the strain carrying the transgene of interest *vsIs33 [dop-3p::RFP]*. *(B)* GFP expression in the genetic balancer strains carrying *myo-2::GFP* markers. *(C)* Schematic of the linkage test. *(D)* Linkage of *vsIs33* over chromosomes, quantified by Linkage Indices shown in a heatmap. *x* axis: indices of the chromosomes; *y* axis: list of the genetic balancers.

We evaluated the linkage of *vsIs33* to autosomes by setting up crosses between our strain of interest (LX811) and a set of balancer strains. These balancer strains are labeled by *myo-2::GFP* markers and have a Green phenotype (GFP expressed in pharyngeal muscles) (Fig. 4*B*). We used the reciprocal translocations *hT2* (34) *[qIs48] (I;III)*, *eT1* (35, 36) *[umnIs12] (III;V)*, and *nT1* (37–39) *[qIs51] (IV;V)* to test for linkage to large portions of chromosomes I and III, III and V, and IV and V, respectively; we also used the inversion *mIn1* (40) *[mIs14] (II)*, to test for linkage to chromosome II (Fig. 4*C* and *SI Appendix*, Extended Methods and Figs. S8-11).

WormPicker picked L4 Green males from the balancer strains and hermaphrodites from the strain of interest (LX811) for mating. WormPicker then screened for F1 Red-Green hermaphrodites, a few of which were subsequently singled. The double-fluorescent F1 hermaphrodites self-fertilized and produced F2s, where the percentages of Red among non-Green animals were assessed to test for linkage between *vsIs33* and particular autosomes. According to classic genetic theory, linkage of the transgene to the tested balancer chromosomes would yield 100% of non-Green animals to be Red, whereas non-linkage of the transgene to the balancer would be reflected by 75% Red among the non-Green progeny.

For testing linkage to the X chromosome, WormPicker generated a genetic cross between LX811 hermaphrodites and wild-type males harboring a dominant extrachromosomal transgene *qnEx615 [myo-2p::GFP]* (NQ1155) (*SI Appendix*, Extended Methods), which helped us to identify F1 cross-progeny. As shown in Fig. 4*C*, F1 Red-Green males were picked out for mating with N2 (wild-type) hermaphrodites. F2s were screened by the robotic system, and the observed percentage of Red among males was used for quantifying the degree to which *vsIs33* is linked to the X chromosome. The theory predicts that none of the F2 males would be Red if X-linked, and that 50% of the F2 males would be Red if not X-linked (*SI Appendix*, Fig. S12).

To quantify the strength of linkage we defined a Linkage Index ranging from -1 to 1, where 1 implies the strongest possible linkage and 0 the weakest (*SI Appendix*, Extended Methods). Our data (Fig. 4*D*) indicate strong linkage of *vsIs33* both to *eT1* and *nT1*, and no linkage to *hT2*, to *mIn1*, or to the X chromosome. Since both *eT1* and *nT1* balance a large portion of chromosome V, our results imply that the RFP transgene is located on chromosome V.

### Genomic integration of a transgene

Transgenic *C. elegans* are usually generated by microinjection, through which cloned DNAs can be delivered to the distal gonadal arm, forming extrachromosomal transgenic arrays (41, 42). To provide stable inheritance and expression, the transgenic arrays can be integrated to the genome. A proven method for genomic integration is to irradiate the strain of interest, causing breakage in chromosomes, which triggers DNA repair, a process through which the transgenic arrays can be ligated to the chromosomes by chance (41). Due to its low frequency, identifying animals with the transgene integrated requires isolating at least several hundred individual worms and screening for 100% inheritance of the transgene in subsequent generations, a highly labor-intensive task (43). In particular, the need to single hundreds of animals can consume a substantial amount of time, even for experienced *C. elegans* researchers.

Using WormPicker, we performed a genomic integration of an *acr-2p::DOP-3* extrachromosomal array labeled by a green fluorescent marker *unc-47p::GFP* (YX293) (Fig. 5). The strain displayed approximately 65% transmission of the transgenic arrays to the next generation. To create the P0 worms, we irradiated 68 L4s with ultraviolet light (254 nm, 30 mJ/cm^2^). WormPicker isolated 216 F1s to individual plates. From these plates, the automated system isolated 612 of their progenies (F2s) to individual plates. The F2 animals were manually screened for 100% transmission of the transgene to F3s and F4s. Among these, two lines were identified as potential independent integrants. For both lines, 100% transmission of the transgene was confirmed over several subsequent generations, consistent with successful array integration.

## Discussion

In this work, we have demonstrated that WormPicker can automate a variety of *C. elegans* genetics procedures usually performed by manual methods. In addition, our user-friendly scripting tools provide flexibility for carrying out customized experiments as well as integrating the system into diverse genetic screens and analyses, potentially including pharmacological screening (44, 45), screening for aging phenotypes (4, 46, 47), and studies of natural genetic variation (48).

Robotic manipulation of *C. elegans* opens possibilities for experiments that would be difficult or impractical for manual methods, especially for those involving a large number of strains or conditions. For example, our laboratory is using this machine to perform a genetic screen for modifiers of stress-induced sleep (49).

The deep-learning-aided machine vision methods we developed here, capable of segmenting individual animals and assaying them across different attributes, may find other applications for *C. elegans* studies, for example in analyses of locomotion (50, 51), aging (52, 53), and sleep (54).

Our self-sterilizing loop design can be used for automatically manipulating other microscopic organisms, for example other nematodes, *Drosophila* larvae, bacteria, and fungi.

Our WormPicker system is primarily made from off-the-shelf components. We have provided a component list including estimated material costs (*SI Appendix*, Table S3) and design files for the custom-made parts (*SI Appendix*, Design File S1). The system control software is accessible at an online repository (55).

It Is important to acknowledge the limitations of our work. For a simple fluorescent animal sorting task, the throughput achieved by WormPicker with a single robotic arm was somewhat smaller than the average of experienced researchers. However, it has the advantage of being able to work continuously without fatigue and to conduct a large number of operations in parallel.

Animals sometimes cluster together in a way that makes it difficult to pick a single individual. We have demonstrated that WormPicker’s machine vision is able to segment individual animals in high-resolution images on highly populated plates (*SI Appendix*, Fig. S6*D*). For the experiments presented here, we programmed the robot to perform intermediate picking (*SI Appendix*, Extended Methods) before transferring animals to their destinations, as a method of handling clusters of worms. In addition, brief blue light illumination (52, 56) or plate vibration could be used to disperse clusters of animals.

Although WormPicker’s machine vision system works well for recognizing and tracking the wild-type animals and mutants tested here, some modifications of our algorithms may be necessary for some mutants with unusual morphologies and behaviors.

We expect all animal manipulations will continue to involve some degree of manual work in conjunction with the automated system. The researcher must prepare plates, load and unload them on the trays, and select the appropriate operations. The effective integration of WormPicker to lab work will require determining which genetic operations are most amenable to automation and which are better left to manual work. The goal of our work is not to fully automate experiments, in the sense that the human is no longer needed, but rather to provide tools to increase the productivity of the researcher.

## Materials and Methods

### Mechanical setup

WormPicker is based on a 1.5 m x 1 m rectangular framing system constructed from aluminum extrusion (OpenBuilds V-Slot 20 mm x 20 mm, 20 mm x 40 mm, and C-Beam 40 mm x 80 mm).

WormPicker’s imaging system and picking arm are moved by a motorized stage along 3 axes: X (80 cm travel), Y (125 cm), and Z (30 cm), In addition, there is a linear carriage under the plate tray that moves the illumination and plate tracking system. All axes are driven by stepper motors (NEMA 23, 1.8 degree per step). For the X and Y axes, motors mounted to the carriage drive motion via a belt attached to both ends of the rail. For the Z axis, a stepper motor drives motion through rotation of a lead screw.

The maximum speed of the X axis was set to 146.67 mm/s, Y axis 146.67 mm/s, and Z axis 8.33 mm/s. The maximum acceleration of the X axis was set to 120 mm/s^2^, Y axis 120 mm/s^2^, and Z axis 10 mm/s^2^. Stepper motors are controlled through a PC via an OpenBuilds BlackBox motion control system using GRBL firmware.

All aspects of the robotic system were controlled by an Origin PC with an Intel(R) Core i9-10900K CPU at 3.7 GHz and 64 GB of RAM, running Windows 10.

An overview of WormPicker hardware architecture is given in *SI Appendix*, Fig. S1.

### Dual-magnification multimodal optical imaging system

To implement bright field imaging, we constructed an illumination system under the platform, in which light from a white LED is diffused by a ground glass and approximately collimated by a Fresnel lens (*SI Appendix*, Fig. S3*A*). The low and high-magnification imaging paths share a common objective lens (achromatic doublet, focal length 100 mm). Light is collected by the objective lens then divided into the low and the high-magnification imaging streams by a beamsplitter (ratio of transmission : reflection is 90% : 10%). The low-magnification image is relayed to a machine vision camera by a camera lens, while the high-magnification image is formed at a CMOS camera through a set of teleconverter lenses and a tube lens. The high-magnification pathway is an infinity-corrected microscopy system that can support both bright field and fluorescence imaging.

For fluorescence imaging (*SI Appendix*, Fig. S3*B* and *C*), two collimated excitation LEDs (center wavelengths 470 nm and 565 nm) and a set of dual-band fluorescence optics (Chroma 59022) were built in the infinity space in the high-magnification pathway. The spectral characteristics of the fluorescence optics were selected to enable both GFP and RFP imaging.

Switching between the imaging modes can be achieved by digital relay circuits controlling the excitation LEDs and the under-platform illumination (*SI Appendix*, Fig. S3*A-C* and Movie S1).

### Robotic picking arm

We built a robotic arm (*SI Appendix*, Fig. S3*F*) for manipulating *C. elegans* on agar media using a wire loop. The robotic picking assembly consists of a linear actuator, three servo motors, and a 3D-printed worm pick (Fig. 1*C*). The linear actuator fine tunes the *z* height of the picking arm through a linear carriage. The servo motors are chained orthogonally to provide 3 degrees of freedom in rotation (θ, ω, φ) (Fig. 1*A*) for the worm pick. The 3D-printed worm pick is mounted to the Servo 1. The design allows two copper wires to fit into the pick stem from the proximal end, and these two wires are connected by a portion of looped platinum wire (90% Pt, 10% Ir, 254 µm diameter) at the distal end of the pick (Fig. 1*C*). The platinum wire is crimped to copper contact pins for attaching to the end of the pick.

To sterilize the pick, the platinum wire loop is connected to a 5 V DC power supply (*SI Appendix*, Fig. S3*G*). The resulting 4.5 A DC current sent through the wire heats the loop to a temperature we estimate to exceed 1,000 °C based on its color (Fig. 1*C* and *SI Appendix*, Movie S3).

We used the wire loop as a capacitive sensing probe (*SI Appendix*, Fig. S3*G*) to monitor contact between the wire and the agar surface, and to provide feedback for picking trajectories. For the sensing circuit to function properly, the platinum wire loop is first disconnected from the heating circuit. The capacitance change due to the contact is sensed by a capacitive touch sensor (SparkFun, AT42QT1011), the voltage output of which is monitored by a data acquisition device (LabJack).

### Pick motion trajectories for manipulating *C. elegans*

To pick up an animal (Fig. 1*D*), the wire pick is positioned above the agar, with a *y*-offset to the target worm (IP). The linear actuator vertically lowers (phase i, change in *z*) the pick until contacting the agar surface (TP) as perceived by the capacitive touch sensor. Next, Servo 2 horizontally swipes (phase ii, change in ω) the pick on the agar surface. Next, Servo 1 and 3 act simultaneously to perform a curved motion (phase iii, change in θ and φ) for picking up the target animal using the outer side of the wire loop (FP). During phases i-iii, the motor speeds are maximized.

To put down the animal (Fig. 1*E*), the pick is positioned above the agar surface (IP). The linear actuator lowers (phase i, change in *z*) the pick until touching (TP), monitored by the capacitive touch sensor. Next, Servo 2 horizontally swipes (phase ii, change in ω) the pick on the agar surface, during which the worm detaches from the wire; for some cases the animal stick to the inner side of the wire loop, the moving stage swipes (phase iii) the pick to −*x*. Finally, Servo 1 raises (phase iv, change in φ) the pick from the agar (FP). The motor speeds are tuned down in different phases, and waiting times were added to phase transitions.

Representative trajectory parameters are listed in *SI Appendix*, Table S5, with the sign convention depicted in Fig. 1*A*. These parameters can be adjusted for different types of animals and scenarios.

### Measurement of viability after automated and manual picking

We manually measured animals’ viability 24 hours after robotic and manual picking. We classified the animals into three categories: alive, dead, and escaped. An animal was classified as alive, if it moved actively; dead, if it lost mobility; and escaped if we could not find it on the plate. For the paralyzed mutant *unc-13* (CB1091), we determined viability by observing pharyngeal pumping.

### Measurement of success rates for the pick-up and put-down procedures

To measure how effectively the system picks up and puts down animals, we manually verified the success of individual pick-up and put-down attempts through the live low-magnification image stream. For accurate manual verification, we limited the number of animals to less than 30 per plate to reduce the probability of picking from and putting down to crowded areas. A pick-up attempt was deemed successful if the animal disappeared from the FOV; failed, if the animal remained in the FOV; and vice versa for verifying a put-down attempt. The success rates of pick-up and put-down attempts were calculated by the number of successes divided by the number of attempts made.

### WormPickerControl

For WormPicker to perform useful work, the basic elements of identifying and transferring animals need to be combined to form multi-step genetic procedures. For this purpose, we developed system control software, WormPickerControl (*SI Appendix*, Fig. S7A). This software is written in Python for the frontend and C++ for the backend.

The frontend is an API through which the user writes scripts to specify tasks to be performed by WormPicker. We developed utilities in the API for initiating resources, generating scripts, managing a set of scripts, and sending the commands to the backend.

The backend contains a library of source scripts, WormPickerLib, each of which enables the hardware to carry out specific tasks. The user can access WormPickerLib through the API and has the flexibility to combine a set of scripts for generating custom protocols. To further improve the speed, we set up multiple threads in the backend for separately handling image acquisition, image processing, hardware control, and script execution.

WormPickerControl contains a database, storing information in CSV format, through which the automated system is able to catalog a set of plates.

We constructed a Mask-RCNN segmentation server, written in Python, responsible for segmenting images acquired from different cameras. The server imports Mask-RCNN models from multiple locally saved PTH files. We built client-server sockets connecting WormPickerLib to the segmentation server, through which images acquired by the hardware are sent to the corresponding Mask-RCNN models, and the inference results are sent back to the software for subsequent processing.

### WormPickerLib

WormPickerLib is a library containing source scripts for performing various procedures, ranging from simple to complex. According to the complexity, elements in the library are categorized into low, mid, and high-level scripts. *SI Appendix*, Table S6 contains brief descriptions for each script in the library.

The low-level scripts enable the system to execute basic actions, such as sterilizing the pick, finding a worm with some desired phenotypes, picking, and putting down the animal. The mid-level scripts are composed of multiple low-level scripts chained in series, e.g., scripts to pick multiple animals with some desired phenotypes from a source to a destination. The high-level scripts are one iteration above the mid-level scripts, enabling the system to carry out complete *C. elegans* genetic procedures. The high-level scripts consist of a group of mid-level scripts arranged in a timed and conditional manner. The user has the flexibility to develop custom procedures using the elements in WormPickerLib.

Taking the genetic mapping experiment (Fig. 4*C*) as an example, *SI Appendix*, Fig. S7B depicts the representative scripts, along with their key input arguments, used for generating a genetic cross between two strains JK2810 and LX811. We wrote high-level scripts (CrossWorms, SingleWorm, and ScreenPlates) for instructing WormPicker to set up the mating of P0, singling F1s, and screening F2s. Execution of CrossWorms relied on calling the mid-level script PickNWorms twice, each time for picking a certain number of males or hermaphrodites to a plate. Then, each time PickNWorms was executed by iterating a set of low-level scripts, combining basic actions of imaging and manipulation.

### Strains

We cultivated all the *C. elegans* strains used in this study at 20°C on nematode growth medium (NGM) plates with OP50 bacteria using standard methods (57). The strains used are described in *SI Appendix*, Table S7. All experiments were carried out at room temperature.

### Data, materials, and software availability

Source data for this paper are available in *SI Appendix*, Dataset S1. Strains used in this study, as tabulated in *SI Appendix*, Table S7, will be available upon reasonable request. Details of WormPicker key components can be found in *SI Appendix*, Table S3. Design files for WormPicker hardware are available in *SI Appendix*, Design File S1. Source code for WormPickerControl, network files for the machine learning algorithms, example scripts for the procedures described in this work, and data processing scripts for the genetic mapping experiment (*SI Appendix*, Fig. S13) can be found on GitHub (55).

## Supporting information

Design File S1

## Acknowledgments

Some strains were provided by the *Caenorhabditis* Genetics Center, which is funded by NIH Office of Research Infrastructure Programs (P40 OD010440). We thank Andrew Ruba, Patrick McClanahan, Michael Hart, and Meera Sundaram for helpful discussions and suggestions. We thank volunteers from the Penn *C. elegans* research community for their participation in the manual picking throughput experiment. This work was supported by the National Institutes of Health (R01NS115995).

## Supporting Information for

### Extended Methods

#### Plate tray platform

We constructed a plate tray platform for housing an array of agar plates. The platform holds eight slide-in trays (Fig. S2*A*), each of which is made of two laser-cut transparent acrylic boards (upper layer 3 mm thick, lower layer 4.5 mm thick) sandwiching a 16.26 µm thick aluminum foil layer (Fig. S2*B*). The function of the foil layer is to increase the capacitance change detected by the capacitive touch sensing circuit (Fig. S3*G*) when the picking wire loop contacts the agar surface. Each tray is held by framing rails, enabling the user to remove or replace the tray by sliding in and out (Fig. S2*A*). Each tray is locked into place by a turn latch (McMaster-Carr 1579N12). Coordinates of individual wells on the platform are calibrated and stored in a configuration file.

#### Plate tracking system

We developed methods for tracking a set of plates based on their barcode labels (Fig. S2*C*). The label contains a machine-readable barcode and human-readable text and is attached to the side of the plate (Fig. S2*D*).

We designed the barcode label to contain 24 black bars, each representing one bit by its thickness (Fig. S2*C*): a thick bar encodes 1, and a thin one 0. Starting from the left, the first 20 bits encode a plate ID number as a binary integer; the next 3 bits encode the MOD-8 checksum of the ID number, used by the algorithm to verify its decoding result. The final (24^th^) bar is always thin, encoding a stop bit.

We built a machine vision module for imaging the barcode from beneath the plate tray (Fig. S2*D*). The barcode label is illuminated by LEDs and a camera captures the image (Fig. S2*E* and *F*).

We developed algorithms (Fig. S2*G-I*) for reading the barcode identifier. To recognize the barcode, we trained a Barcode-finding Mask Regional Convolutional Neural Network (1) (BcMRC). We curated a training dataset containing 520 images (648 pixels x 486 pixels size and 5.8 cm x 4.4 cm FOV) in which the contours of individual barcodes were manually labeled. BcMRC was adapted from a pretrained ResNet-50-FPN backbone (1). The training was performed with a Stochastic Gradient Descent (SGD) optimizer and the model converged after 9 epochs. The model generates a contour segmentation for the object most likely to be the barcode in the image (Fig. S2*G*) (we assume there is only one barcode visible in the image). The mean Intersection over Union (IoU) score of the mask prediction with respect to the ground truth is 0.942 (N = 80).

We generate a scanning line running through the BcMRC-predicted mask (Fig. S2*G*). Intensities of some pixels above and below the scanning line are averaged for calculating the intensity profile (Fig. S2*H*). Then, the intensity profile is binarized by a threshold *H* (Fig. S2*I*), *H* = (*p*_0.1_ + *p*_0.9_)⁄2, where *p*_0.1_ and *p*_0.9_ are the 10^th^ and 90^th^ percentile intensity values. After the thresholding, a bit is decoded as 1 if the width of the black bar is larger than its complementary white bar right next to it, and 0 otherwise (Fig. S2I). The decoded ID number is verified by its checksum. An error is reported if the verification fails.

Individual plates are cataloged in a database (Fig. S7*A*) based on their ID numbers. Information stored in the database includes but is not necessarily limited to the positions on the tray, strain names, genotypes, phenotypes, and histories. The database is arranged in a human and machine-readable CSV format.

#### Image focus control

The distance between the main objective lens and the agar surface is monitored by a custom-built laser-based level monitor (Fig. S3*D*). A 532 nm laser containing a cylindrical lens generates a collimated laser line lying in the *y* axis. The laser line is focused by a lens (focal length 100 mm) and projected onto the agar surface at an oblique angle (Fig. S3*D*). Then, the distance between the objective lens and the agar surface can be monitored by the position of the laser line observed by the imaging system. The 3D motorized stage brings the objective to the correct position, at which the agar surface lines up with its focal plane, by aligning the observed laser line to a previously calibrated position.

#### Lid handler

Lids are manipulated by two custom-built vacuum grabbers mounted to the 3D motorized stage (Fig. S3*E*). Each grabber consists of a pair of 3D-printed sockets holding a vacuum tube, motorized by a linear actuator (Actuonix, PQ12-30-12-S). A tube fitting is attached to the end of the vacuum line. The height of the vacuum line is controlled by the linear actuator and the vacuum is switched on or off by a solenoid valve (Granzow, H2U29-00Y).

To remove a lid, the main moving gantry positions the vacuum holder above the lid, the solenoid valve activates the vacuum, the linear actuator lowers the vacuum holder, the lid becomes attached by suction to the holder, and the linear actuator raises the holder and the lid (Movie S2). To restore a lid, this process is reversed.

#### Pick focus control

To bring the pick tip to the optimal focus in the high-magnification FOV, the system monitors the mean intensity of the top 20% darkest pixels 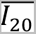 of the image as a measure of focus (Fig. S4). The system sets the height *z*_*f*_ at which the 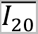 is minimized as the focus height of the pick tip. The system records the positions read out from the linear actuator and the three servos, i.e., (*l_f_*, *s*_1*f*_, *s*_2*f*_, *s*_3*f*_), when the pick tip is at the focus position (*x_f_*, *y_f_*, *z_f_*).

To pick a worm, the system positions the pick tip right above the remembered focus position, i.e., (*x_f_*, *y_f_*, *z_f_* + Δ*z*), where Δ*z* is a predefined small amount of height offset (approximately 2 mm). We set the position (*x_f_*, *y_f_*, *z_f_* + Δ*z*) to be the initial position (IP in *Main Manuscript*, Fig. 1*D* and *E*), prior to carefully lowering the pick until touching the agar.

We measured the precision of moving the pick tip to the remembered focus position in the *x* and *y* directions. The mean differences between the actual position and the focus position are 290 µm (SD = 161 µm) in *x* and 250 µm (SD = 173 µm) in *y* (N = 342). This precision allows the system to move the pick to the focus position directly in a feed-forward manner, helping to improve the worm picking speed.

#### Machine vision

##### *C. elegans* tracking in the low-magnification imaging stream

We used a combination of machine learning and motion detection for tracking *C. elegans* over the plate in the low-magnification imaging stream.

We trained WormCNN, a convolutional neural network (CNN) for making a binary prediction of whether a 1.58 mm x 1.58 mm region of interest (ROI) contains one or more worms. The structure of the WormCNN is shown in Fig. S5*A*. We curated a training dataset containing 43,568 ROI images (20 pixels x 20 pixels size), 50% worms, and 50% backgrounds (Fig. S5*B*). With a batch size of 24 and an SGD optimizer (momentum = 0.9), the model converged after 10 training epochs. The accuracy of the network is 97.23% on 10,784 test images (balanced test set, i.e., half worms and half backgrounds).

In addition to the results from the machine learning, we constructed a simple motion detector for identifying animals over the plate. The entire FOV is divided into multiple 0.63 mm x 0.63 mm ROIs. We took two frames Δ*t* seconds apart, subtracted the pixel values, and summed up the pixel differences Δ*p* within the ROI. If the Δ*p* is higher than a threshold *p_n_*, i.e., Δ*p* > *p_n_*, we consider the ROI containing animals. Values of Δ*t* and Δ*p* were calibrated on the animals of interest for the optimal performance.

Combining WormCNN and motion detection, the machine vision for *C. elegans* tracking functions in two modes: Finding mode and Tracking mode.

To rapidly identify animals over the plate, the Finding mode runs the motion detector over the entire FOV. Then it feeds the ROIs with the movement detected to WormCNN to check whether any animals are present in these ROIs. If so, the system sets the centroid coordinates of the ROIs to be the coordinates of the animals identified.

The Tracking mode tracks the animals using WormCNN, supplemented by subpixel processing. For an animal having a known coordinate *C*_*n*−1_ = (*X*_*n*−1_, *Y*_*n*−1_) for the frame *n* − 1, the Tracking mode updates its coordinate for the frame *n* by a few steps.

First, it constructs a 2.53 mm x 2.53 mm rectangular tracking region, centering at *C*_*n*−1_. Second, it splits the tracking region into multiple 1.58 mm x 1.58 mm ROIs separated by a constant step size (0.39 mm) in *x* and *y* directions. Each ROI is sent to WormCNN for generating a confidence score *cs*. The confidence score *cs* is the output of the final fully connected layer of WormCNN, indicating the likelihood of the input ROI containing any worms. Third, it calculates the centroid of the *cs* for all the ROIs in the tracking region and sets it to be the animal’s coordinate in the frame *n*, i.e., *C*_*n*_ = (*X*_*n*_, *Y*_*n*_). *X*_*n*_ and *Y*_*n*_ are given by

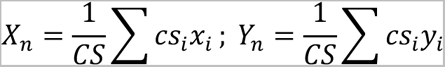

where *cs_i_* is the confidence score of the *i*-th ROI; *x_i_*, and *y_i_* the *x* and *y* coordinates of the centroid of the *i*-th ROI; *CS* the sum of the confidence score for all the ROIs, i.e., *CS* = ∑ *cs_i_*.

In the equation above, *x_i_*, and *y_i_* are dependent on the coordinate of the animal in the frame *n* − 1, i.e., *C*_*n*−1_, the last known coordinate of the animal being tracked. Therefore, *x_i_* and *y_i_* can be written as *x*_*n*\*1,+_ and *y*_*n*\*1,+_ for indicating the temporal dependency. In short, the Tracking mode updates the coordinate of the animal for the current frame (*X*_*n*_, *Y*_*n*_) by its last seen coordinate (*X*_*n*−1_, *Y*_*n*−1_) by

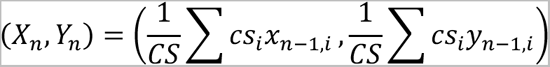

Using the Finding and the Tracking mode, WormPicker rapidly recognizes and moves individual animals to the high-magnification FOV, preparing for imaging and manipulation. This process involves a few steps.

First, the system runs the Finding mode to obtain the coordinate for a worm (*X*_*W*_, *Y*_*W*_). Then, the main gantry moves the image of the animal to the high-magnification FOV by translating by Δ*X*_g_ and Δ*Y*_g_ in the *x* and *y* directions. Δ*X*_g_ and Δ*Y*_g_ are given by

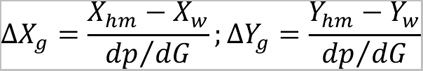

where *X_h_*_*m*_ and *Y_h_*_*m*_ are the *x* and *y* coordinates of the high-magnification FOV; *dp*⁄*dG* the amount of the pixel shift corresponding to one unit movement in the main gantry. By making the gantry movement above, the animal is viewed in proximity to the high-magnification FOV.

The freely roaming animal can be kept centered at the high-magnification FOV by running the Tracking mode while the gantry is continuously making small-step jogging motions in *x* and *y*. The movements of the jogging in *x* and *y* are given by

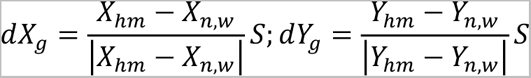

where *X*_*n*,_*_w_* and *Y*_*n*,_*_w_* are the *x* and *y* coordinates of the worm in the current frame *n*, and *S* the predefined step size for the gantry jogging.

##### Pick tracking in the low-magnification imaging stream

To track the pick wire loop in the low-magnification imaging stream, we trained PickCNN, a CNN for determining whether a 1.26 mm x 1.26 mm ROI contains the pick tip. The backbone of PickCNN is the same as WormCNN (Fig. S5*A*). We curated a training dataset containing 872,550 ROI images (20 pixels x 20 pixels size), 50% containing the pick tip and 50% the backgrounds (Fig. S5*C*). The network was trained on a GPU (NVIDIA GeForce GTX 1660) using an SGD optimizer. The accuracy of the model is 99.36% on 50,400 test images (balanced test set, i.e., half pick tips and half backgrounds).

We applied PickCNN, supplemented with subpixel processing, to track the pick tip in real time. A 20 mm x 15 mm rectangular region centered at the low-magnification FOV is delineated for tracking the pick tip inside the region. This tracking region is split into multiple 1.26 mm x 1.26 mm ROIs with a constant skip step (0.32 mm) in *x* and *y* directions. Each ROI is fed to PickCNN for generating a confidence score *cs*. The confidence score *cs* is the output of the final fully connected layer of PickCNN, indicating the likelihood of the ROI containing the pick tip.

Assuming there is only one picking arm visible, among all the ROIs, the one with the highest *cs* is the closest to the actual pick tip. We use (*x_h_*, *y_h_*) to denote the centroid coordinate of the ROI having the highest *cs*. By performing subpixel processing surrounding the (*x_h_*, *y_h_*), the coordinate of the pick tip (*X*_*p*_, *Y*_*p*_) can be inferred by

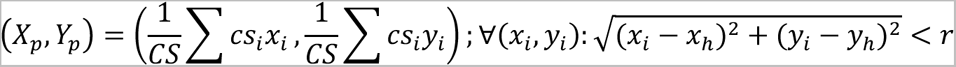

where *cs_i_* is the confidence score of the *i*-th ROI; *x_i_*, and *y_i_* the *x* and *y* coordinates of the centroid of the *i*-th ROI; *CS* the sum of the confidence score for all the ROIs, i.e., *CS* = ∑ *cs_i_*. The equation above indicates that all the ROIs having the distance to (*x_h_*, *y_h_*) less than *r* are taken into the subpixel processing for determining the pick tip coordinate (*X*_*p*_, *Y*_*p*_). We set *r* = 1.89 mm.

We measured the precision of the pick tracking. The mean differences between the tracked position and the actual position are 94.9 µm (SD = 43.0 µm) in *x* and 284.5 µm (SD = 147.2 µm) in *y* (N = 225).

##### *C. elegans* segmentation in the high-magnification imaging stream

We trained a Worm-finding Mask Regional Convolutional Neural Network (1) (WorMRC), for pixel-wise segmentation for individual animals in the high-magnification FOV. We curated a training dataset containing 2,697 images (612 pixels x 512 pixels size and 1.88 mm x 1.57 mm FOV), over different developmental stages, ranging from L1 larvae to adult. For each training image, we manually labeled the contours for individual animals. WorMRC was adapted from a pretrained ResNet-50-FPN (1) backbone, and was trained at a GPU (NVIDIA GeForce GTX 1660 Ti) using an SGD optimizer (momentum = 0.9). The model converged after 10 training epochs.

The network outputs bounding boxes for the objects that are likely to be the worm and generates a contour for the object within the bounding box. WorMRC robustly segments individual animals over complex backgrounds (Fig. S5*D*). We plot the areas of the contours for individual animals (of mixed stages, L1s - adults) predicted by WorMRC against the results from manual segmentation in Fig. S5*E*. The WorMRC predicted contour area presents high consistency with the ground truth, *R*^2^ = 0.972 (N = 690).

##### Developmental stage determination

The developmental stages of the animals in the high-magnification image were inferred by their lengths. WorMRC first generated contours for individual animals. The centerline of the individual contour was obtained by the methods previously reported (2). We used previously measured lengths for worms of various developmental stages (3) to obtain decision boundaries for approximately identifying the developmental stage: L1s are < 370 µm, L2s 370 – 500 µm, L3s 500 – 635 µm, L4s 635 – 920 µm, adults > 920 µm.

##### Morphological analysis

To perform the genetic cross between the *dpy-5* mutant and *dop-1p::GFP* transgenic (*Main Manuscript*, Fig. 3) we first developed an algorithm for identifying the Dumpy (Dpy) phenotype. The *dpy-5* mutant displays a shorter and stouter morphology than control animals. We measured the aspect ratio *AR* of the animal, 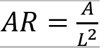, where *A* is the area of the contour segmented by WorMRC and *L* the length of the centerline of the worm contour (2). We used the automated system to record images for *dpy-5* mutants (CB61) and wild-type animals (N2) (N = 190).

Based on this dataset, we determined an *AR*-based threshold for identifying Dpy animals. The values of *AR* for individual Dpy and wild-type animals are plotted in Fig. S5*F*. With an 80%-20% train-test split, we constructed a one-dimensional support vector machine (SVM) to classify the Dpy and wild-type phenotypes based on the *AR*. The SVM-generated decision boundary gives 100% accuracy on both the training and the test sets (Fig. S5*F*). We used this threshold to classify future samples: if a worm has an *AR* greater than the threshold, then the worm is classified as Dpy; otherwise, it is classified as having a wild-type morphology.

##### Sex determination

*C. elegans* males and hermaphrodites display distinct tail morphologies, with a fan structure visible in the tails of males but not hermaphrodites. We trained a Sex-determining Mask Regional Convolutional Neural Network (1) (SexMRC), for determining the sexes of *C. elegans* based on the high-magnification images. We curated a training dataset containing 6,860 images (612 pixels x 512 pixels size and 1.88 mm x 1.57 mm FOV), for which the contours of heads, tails for hermaphrodites, and tails for males, were manually labeled. SexMRC was adapted from a pretrained ResNet-50-FPN (1) model and was trained with an SGD optimizer (momentum = 0.9) using a GPU (NVIDIA GeForce GTX 1660 Ti). The model converged after 7 training epochs. The model outputs bounding boxes for the objects that are likely to be *C. elegans* heads, tails for hermaphrodites, and males, as shown in Fig. S5*G*. The network identifies males with a 100% precision and an 85.71% sensitivity (recall) and identifies hermaphrodites with a 92.45% precision and a 92.45% sensitivity (recall) (N = 74).

##### Embryo detection

We trained an Embryo-finding Mask Regional Convolutional Neural Network (1) (EmbMRC), for identifying embryos in the high-magnification images. We curated a training dataset containing 454 images (612 pixels x 512 pixels size and 1.88 mm x 1.57 mm FOV), for which the contours of unhatched embryos were manually labeled. EmbMRC was adapted from a pretrained ResNet-50-FPN (1) backbone and was trained using an SGD optimizer at a GPU (NVIDIA GeForce GTX 1660 SUPER). The model converged after 10 training epochs. The model outputs bounding boxes for the objects that are likely to be the embryo and generates a contour for the object within the bounding box (Fig. S5*H*). Fig. S5*H* shows representative images along with the segmentation when a few, a medium number of, and a larger number of embryos are present in the FOV. The numbers of embryos counted by EmbMRC are highly consistent with the results from manual counting (*R*^2^ = 0.9772, N = 124) (Fig. S5*I*).

##### Fluorescence phenotyping

For fluorescence phenotyping, the automated system moves to view individual worms in the high-magnification FOV and captures one bright field and one subsequent fluorescence frame. The bright field frame is delivered to WorMRC for obtaining contour segmentations for the animals. The fluorescent frame is binarized by an intensity threshold *h*_*i*_ and the bright spots obtained after the thresholding are filtered according to their sizes, for reducing noise. The bright spots with sizes larger than a threshold ℎ_*s*_ are kept as true fluorescent spots; otherwise, they are discarded. Both *h*_*i*_ and ℎ_*s*_ were calibrated on the fluorescent strains of interest. For each animal identified in the bright field image, we dilated the contour mask to compensate the potential mismatch between the bright field and fluorescent frames caused by the animal’s free-roaming behaviors during the elapsed time (about 250 ms) for the frame acquisition. The animals cut off by the FOV boundaries were filtered out for fluorescence phenotyping, preventing false negatives. The parameters for fluorescence imaging and processing used for our experiments are given in Table S4.

For a moderately bright fluorescent strain (LX811), the algorithm classifies fluorescence phenotypes for individual animals of mixed stages with an accuracy of 97.16% (N = 211).

##### Automated verification of the transfer results

For the system to keep track of the working progress, we programmed the machine vision to verify whether the animal was successfully transferred to the destination. First, the robotic arm attempts to put down the animal in the high-magnification FOV, and it observes whether any worms can be found in the FOV using WorMRC. If so, the transfer is deemed a success; otherwise, continuing with the next step. Second, it observes whether any animals crawl out from the proximity of the spot where the worm was put down via the low-magnification FOV. If there are any such animals, the operation is deemed a success. The automated verification strategy achieves a 97.22% accuracy for verifying the results of the transfer attempts (N = 108).

### Intermediate picking

Depending on the application, the automated system may be required to precisely transfer a single animal with some desired phenotypes and stray animals are strictly prohibited, for example, isolating F2s for identifying a transgenic integrant (*Main Manuscript*, Fig. 5). We programmed the automated system for those experiments to perform intermediate picking if multiple worms or/and unhatched embryos are identified in the high-magnification FOV.

The system first inspects all the animals in the image, if none of them carries the desired phenotypes, the system inspects other animals; if at least one carries the desired phenotypes, the robotic arm picks up the target animal, possibly along with surrounding ones. Then, the system puts the animals down onto a fresh intermediate plate and waits a few seconds allowing the worms to move. The system then picks up the single animal with the desired phenotypes.

### Theory of the genetic mapping experiment

Using the automated methods, we genetically mapped an RFP transgene *vsIs33 [dop-3p::RFP]* using classic genetics. We tested the linkage of *vsIs33* to different chromosomes by setting up genetic crosses between the strain of interest (LX811) and a set of genetic balancers labeled by GFP markers.

The first genetic balancer strain we used was JK2810, carrying a *hT2* (4) reciprocal translocation in I and III, for determining linkage to I and III. As shown in Fig. S8, WormPicker crossed JK2810 males with LX811 hermaphrodites and picked out F1 cross-progenies that were both Red and Green. According to the theory, the F2 self-progenies descending from these double-fluorescent F1s would display a segregation pattern for the Red and the Green, as shown in the Punnett squares (Fig. S8). If *vsIs33* is linked to neither I nor III, the theory predicts 75% of the non-Green animals to be Red, i.e., P (Red | non-Green) = 0.75 (Fig. S8*A*); while if *vsIs33* is linked to either I or III, 100% of the non-Green would be Red, i.e., P (Red | non-Green) = 1 (Fig. S8*B*).

We determined the linkage of *vsIs33* to other autosomes using the same scheme. The automated system generated crosses between LX811 and other GFP-labeled genetic balancers, picked double-fluorescent F1 cross-progeny, and screened for the segregation pattern for the Red and the Green in F2s. These balancer strains include VC170, carrying a *mIn1* (5) inversion in II (Fig. S9), CGC34, carrying an *eT1* (6, 7) reciprocal translocation in III and V (Fig. S10), JK2958, carrying a *nT1* (8–10) reciprocal translocation in IV and V (Fig. S11). For all the autosomal genetic balancers, the theory predicts 75% of the non-Green F2s would be Red if unsuccessfully balanced, and 100% if successfully balanced.

To test the linkage of *vsIs33* to the X chromosome requires a different scheme (Fig. S12). WormPicker crossed wild-type males harboring a dominant extrachromosomal transgene *qnEx615[myo-2p::GFP]* (NQ1155) with LX811 hermaphrodites, and picked out double-fluorescent F1 males, which were subsequently crossed with wild-type (N2) hermaphrodites. The theory predicts that none of F2 males would be Red if the transgene of interest is X-linked, and 50% if not X-linked.

For each linkage test for the autosomes described above, WormPicker screened over the F2s and obtained the numbers of the non-Green animals that were Red, *N*_*A*1_, and non-Red, *N*_*A*2_, and the total numbers of the non-Green animals found, *N* = *N*_*A*1_ + *N*_*A*2_. As shown in Fig. S13*A*, the red bars represent the observed percentages of the Red among the non-Green animals, 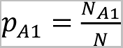, while the gray bars the percentages of the non-Red among the non-Green animals, 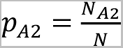. The error bars indicate the standard deviation *SD*_*A*_ of the observed percentages under the autosome-unlinked assumption, where 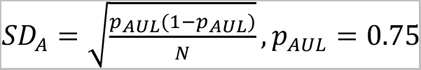.

Similarly, for testing the X-linkage, the automated system screened over the F2s and obtained the number of males that were Red, *N_x_*_1_, and non-Red, *N_x_*_2_, and the total number of males found *N* = *N_x_*_1_ + *N_x_*_2_. As shown in Fig. S13*B*, the red bar represents the observed percentage of Red among males, 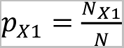, while the gray bar the percentage of non-Red among males, 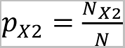. The error bars indicate the standard deviation *SD_x_* of the observed percentages under the X-unlink assumption, where 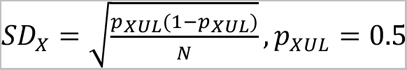.

To evaluate the linkage strength, we developed a Linkage Index to quantify the similarity between the theoretical and the observed link (Fig. S13*C*). To derive the Linkage Index, we first vectorize the link pattern in theory *W*_*L*_, the unlink pattern in theory *W*_*UL*_, and the observed pattern *W_obs_*:

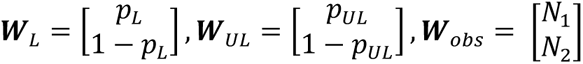

For the autosome-linkage test, *p*_*L*_ and *p*_*UL*_ are the fractions of the non-Green animals that would be Red for the link and the unlink patterns in theory, i.e., *p*_*L*_ = 1, *p*_*UL*_ = 0.75; *N*_1_ and *N*_2_ are the observed count numbers of the non-Green animals that were Red and non-Red. For the X linkage test, *p*_*L*_ and *p*_*UL*_ are the fraction of the males that would be Red for the X-link and X-unlink patterns in theory, i.e., *p*_*L*_ = 0, *p*_*UL*_ = 0.5; *N*_1_ and *N*_2_ are the observed count numbers of males that were Red and non-Red.

Let *e*_*L*_ and *e*_*UL*_ be the unit vectors pointing to the same directions as *W*_*L*_ and *W*_*UL*_, i.e.,

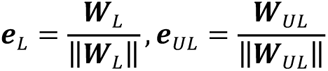

(‖ ‖ denotes the L2-norm of the vector). Then we set *e*_*L*_ and *e*_*UL*_ as the basis vectors and *W*_*obs*_ can be expressed in terms of the basis

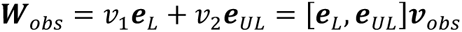

where 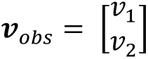 is the coordinate of *W_obs_* with respect to the basis *e_L_* and *e_UL_*.*v*_*obs*_ can be obtained by

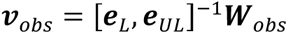

where [ ]^-1^ denotes the matrix inverse. The coordinates of the basis vectors themselves are

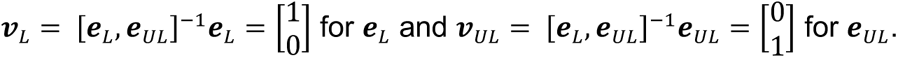

We define the Linkage Index *Lidx* as the cosine similarity between the *v*_*obs*_ and *v*_*L*_:

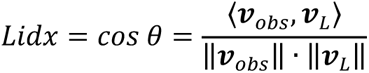

where *θ* is the angle formed by the *v*_*obs*_ and *v*_*L*_ (Fig. S13*C*) and 〈 〉 denotes the vector inner product. If the observed pattern is close to the link pattern, then *θ* → 0, yielding *Lidx* → 1, implying a strong linkage to the chromosomes of interest. If, on the other hand, the observed pattern is similar to the unlink pattern, the *θ* → 90°, yielding *Lidx* → 0, implying a weak linkage. Note that we cannot draw any conclusions when *Lidx* → −1 because it suggests that the observed pattern is far deviated from both the link and the unlink patterns.

The observed Linkage Indices for *vsIs33* over different chromosomes are shown in Fig. S13*D*. We found that *vsIs33* displays strong linkage to either III or V, and to either IV or V, suggesting the transgene of interest is linked to V.

---

**Fig. S1.**
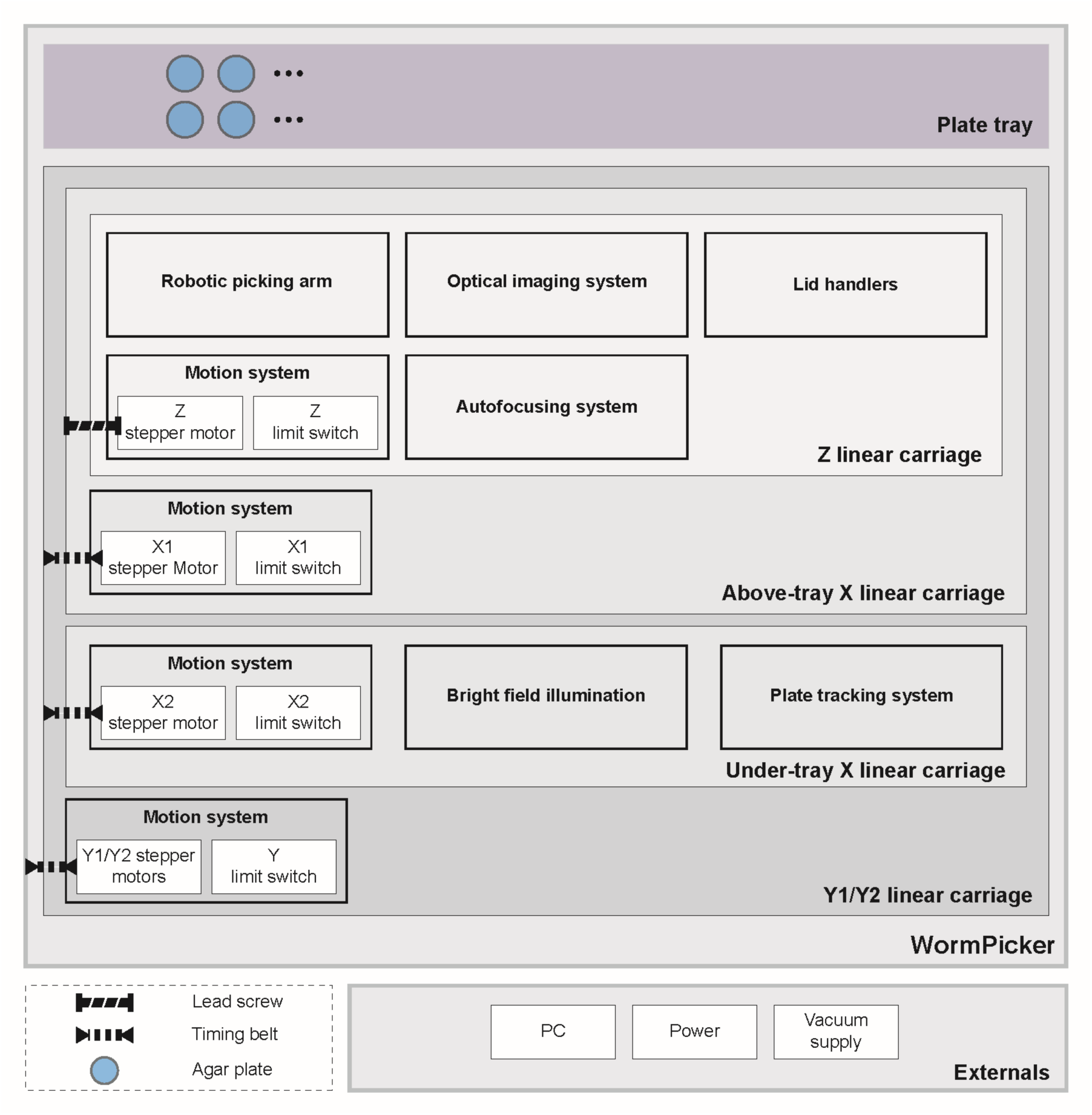
WormPicker hardware architecture. WormPicker is composed of a plate tray platform housing an array of agar substrates and linear carriage assemblies built in Cartesian coordinates, i.e., X, Y, and Z. The Z linear carriage motorizes imaging and manipulation assemblies, including a robotic picking arm, an optical imaging system, two lid handlers, and an image autofocusing system. The under-tray X linear carriage motorizes supplementary tools, including a bright field illumination and a plate tracking module.

**Fig. S2.**
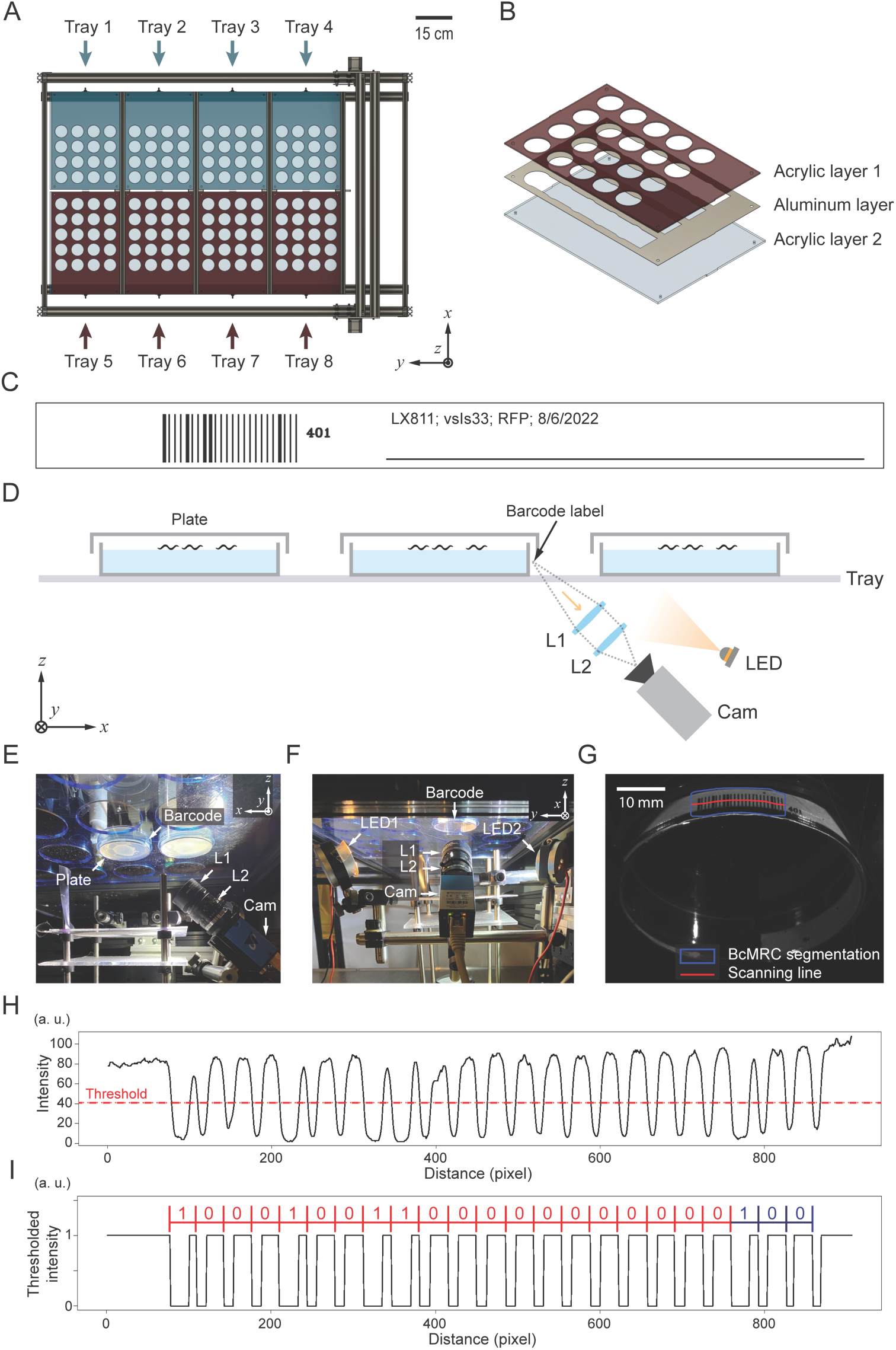
WormPicker plate tray platform and plate tracking assemblies. *(A)* Top view. The platform is composed of 8 easy slide-in plate trays. *(B)* Each slide-in plate tray consists of two transparent laser-cut acrylics sandwiching an aluminum layer. *(C)* A barcode label. *(D)* Machine vision system for barcode reading. *(E* and *F)* Front and side view of the barcode-reading apparatus. *(G)* Camera-captured image of the barcode label with BcMRC segmentation and a scanning line. *(H)* Profile of pixel intensity along the scanning line drawn in *(G)*. Red dashed line: threshold for binarizing the intensity profile. *(I)* The binarized intensity profile of *(H)*. Red: decoded 20-bit number; Blue: 3-bit checksum.

**Fig. S3.**
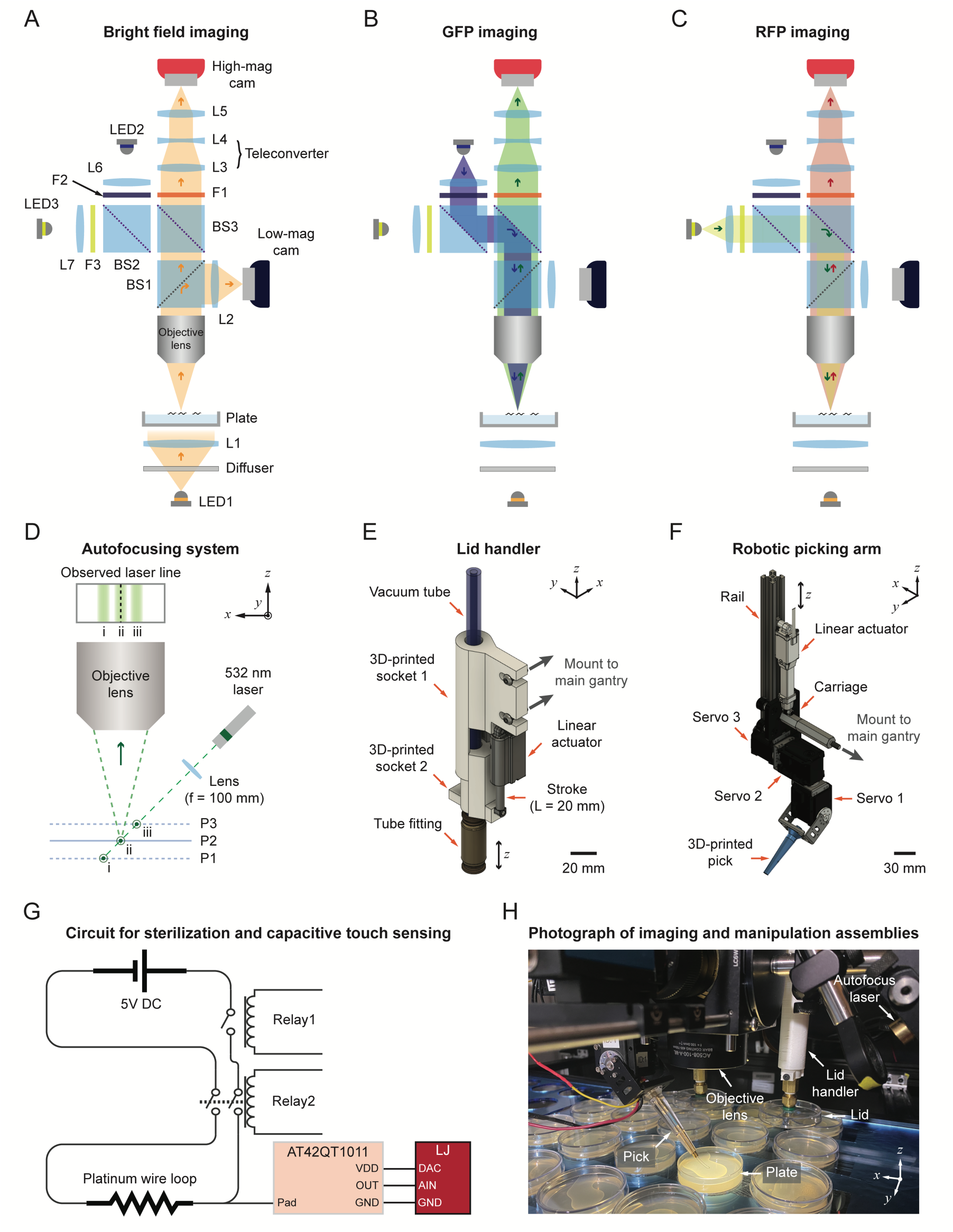
WormPicker imaging and manipulation assemblies. *(A-C)* Schematic of *(A)* bright field, *(B)* GFP, and *(C)* RFP imaging. BS1: beamsplitter (transmission : reflection = 90% : 10%). BS2 and 3: long-pass and dual-band dichroic beamsplitters. F1: dual-band emission filter. F2 and 3: excitation filters for GFP and RFP. L1: Fresnel lens. L2: machine vision lens. L3 and 4: teleconverter lens pair (f = 300 mm and -100 mm). L5: tube lens. L6 and 7: collimating lenses. LED1-3: white, 470 nm, and 565 nm LED. Objective lens: achromatic doublet lens (f = 100 mm). *(D)* Schematic of autofocusing system. P1-3: different positions of an agar surface, where P2 is the focus position. i-iii: positions of a laser line projected to the agar surface at different planes, i.e., P1-3. Dashed line: pre-calibrated optimal position. *(E)* Schematic of lid handler. *(F)* Schematic of robotic picking arm. *(G)* Electric circuit for pick sterilization and capacitive touch sensing. AT42QT1011: capacitive touch sensor; LJ: LabJack data acquisition device. *(H)* Photograph of imaging and manipulation assemblies.

**Fig. S4.**
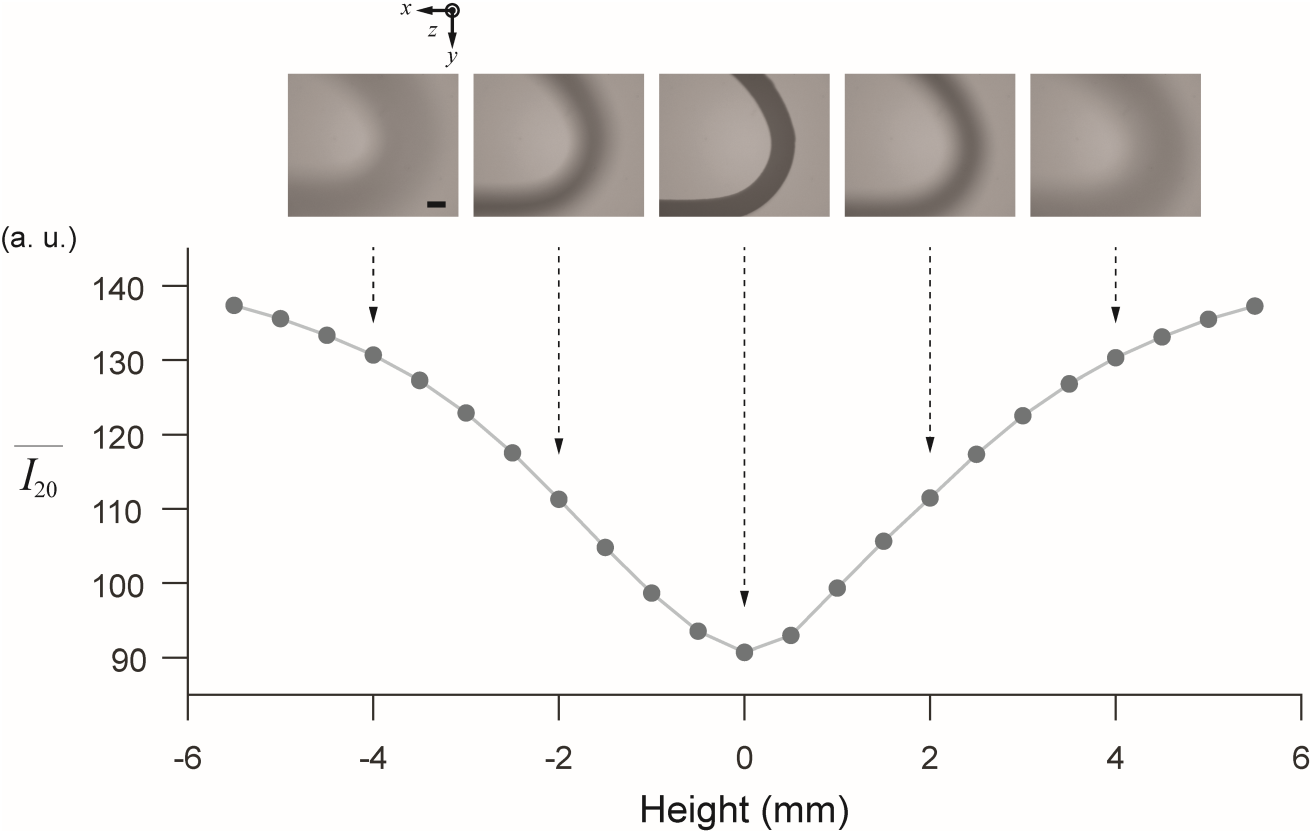
Mean intensity of the top 20% darkest pixels in the high-magnification 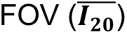 for the pick tip at different heights in ***z***. Above: pick tip images. Scale bar: 200 µm.

**Fig. S5.**
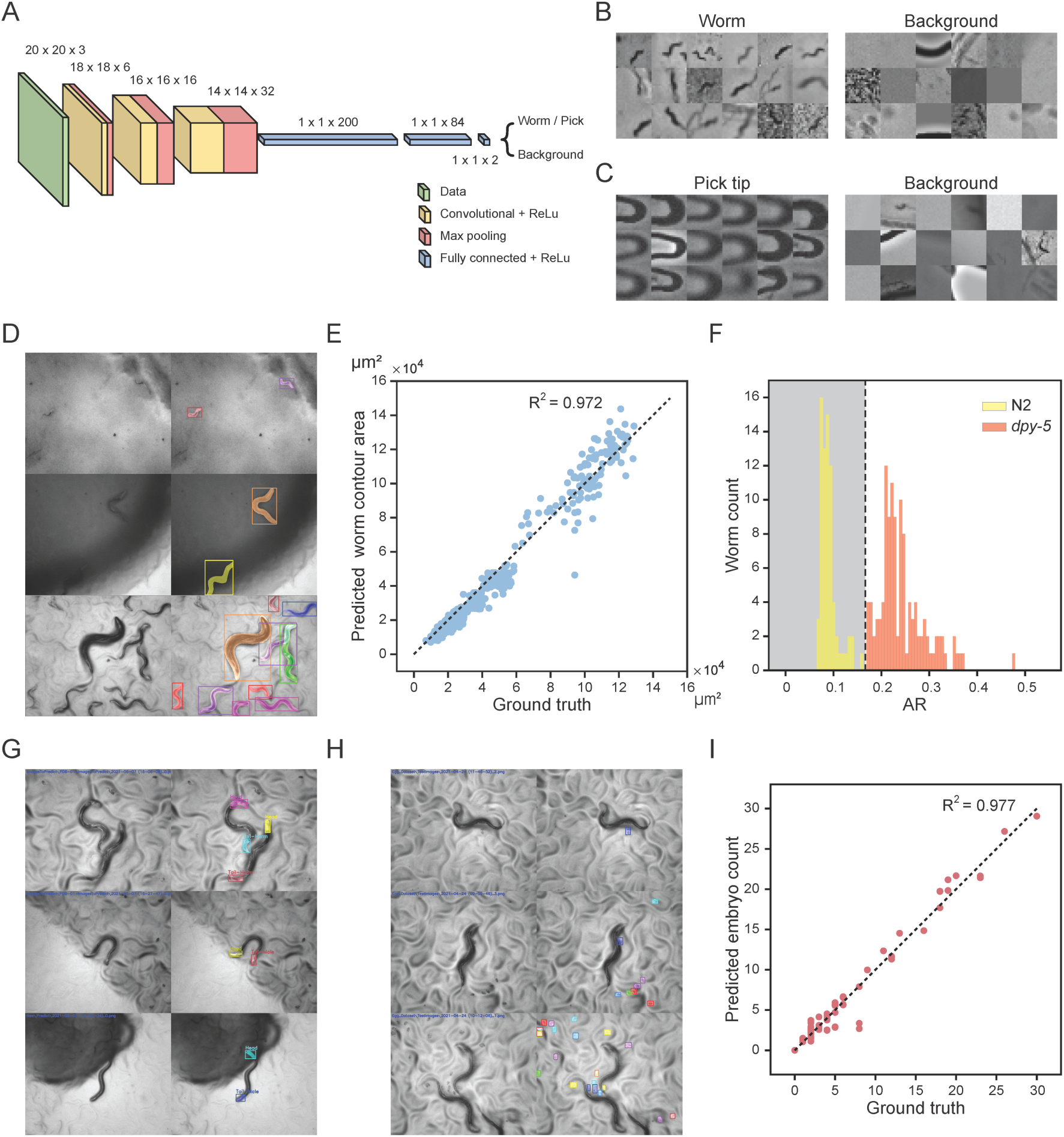
Machine vision analysis. *(A)* Structure of WormCNN and PickCNN. *(B)* Representative images of worm (left) and background (right) for WormCNN training. FOV: 1.58 mm x 1.58 mm. *(C)* Representative images of pick tip (left) and background (right) for PickCNN training. FOV: 1.26 mm x 1.26 mm. *(D)* Worm segmentations by WorMRC (left: raw images; right: segmentations). FOV: 1.88 x 1.57 mm. Individual worms are labeled with different colors. *(E)* Scatter plot of areas of individual worm contours (of mixed stages, L1-Adult) segmented by WorMRC versus ground truth (N = 690). Dashed line: ***y*** = ***x***. *(F)* Histogram of aspect ratios (ARs) of wild-type (N2) and Dpy (*dpy-5*) animals (N = 190). Black dashed line: decision boundary for classifying wild-type (gray region) and dumpy morphology (white region). *(G)* Sex determination by SexMRC (left: raw images; right: segmentations). Individual heads and tails are labeled with different colors. Recognized types of the tails are indicated by the text. *(H)* Embryo detection by EmbMRC (left: raw images; right: segmentations). Individual embryos or clusters of embryos are labeled with different colors. *(I)* Scatter plot of numbers of embryos detected by EmbMRC versus ground truth (N = 124). Dashed line: ***y*** = ***x***.

**Fig. S6.**
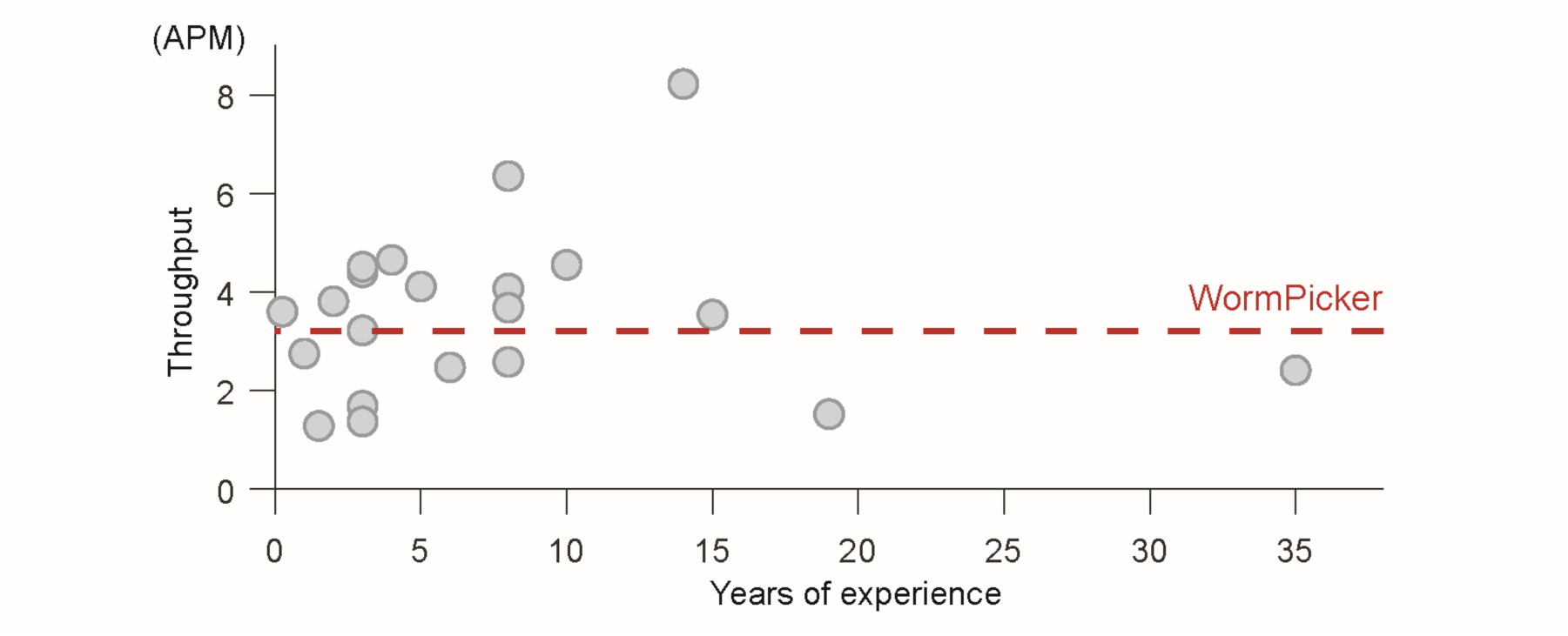
Scatter plot of picking throughput (unit: Animals per Minute, APM) versus years of experience for a fluorescent animal sorting task performed by a group of researchers (N = 21). Mean and median years of experience: 7.61 and 5 years. Mean, median, and standard deviation (SD) of manual throughput: 3.56, 3.60, and 1.67 APM. Dashed line: automated throughput, 3.21 APM (SD = 0.66 APM, N = 38).

**Fig. S7.**
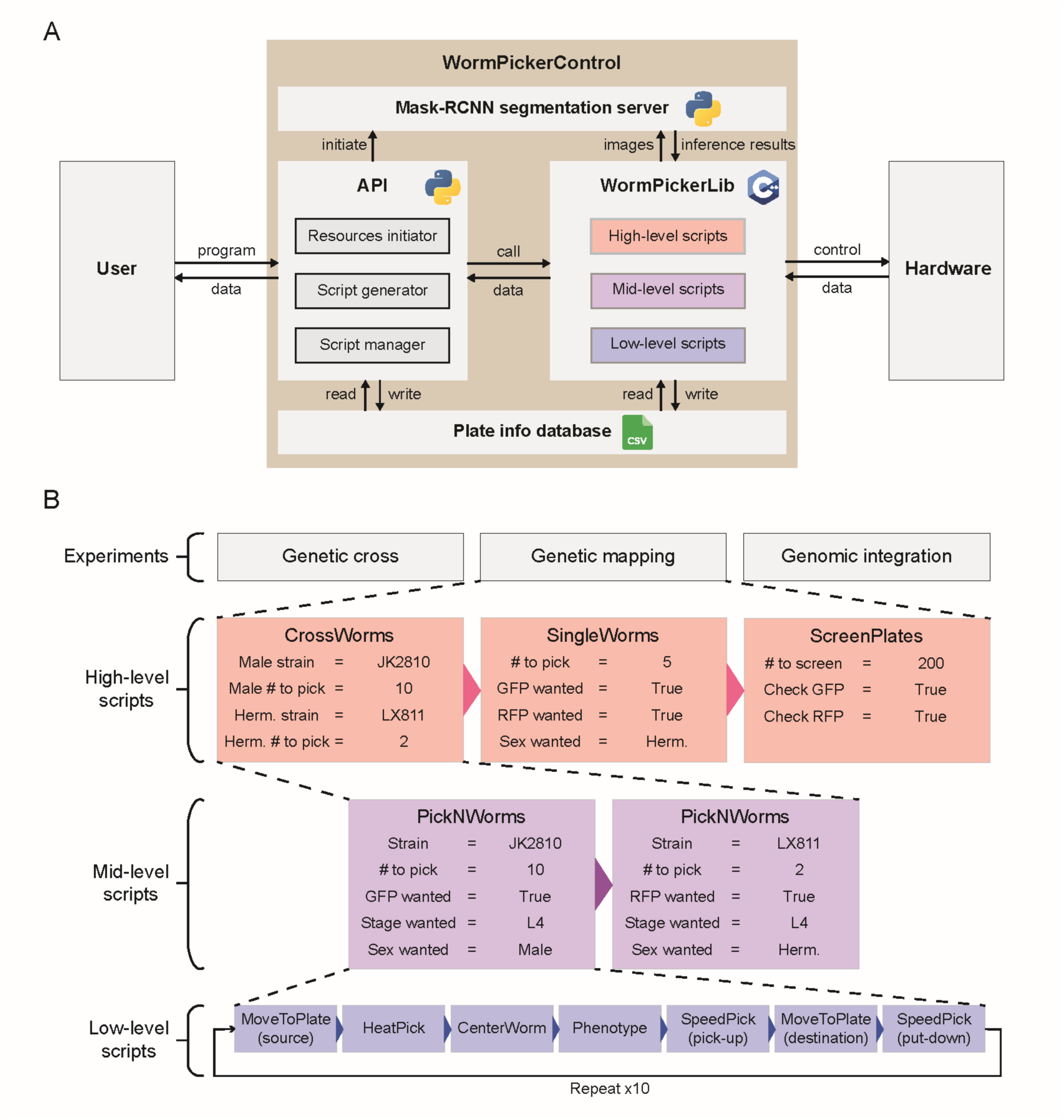
WormPickerControl and WormPickerLib. *(A)* Schematic of WormPickerControl. WormPickerControl is composed of four modules, including an Application Programming Interface (API), WormPickerLib (a library of source scripts), a Mask-RCNN segmentation server, and a database cataloging plate information. *(B)* Schematic of WormPickerLib, demonstrating its hierarchical structure by taking the genetic mapping experiment as an example (cross JK2810 with LX811, as shown in *Main Manuscript*, Fig. 4C).

**Fig. S8.**
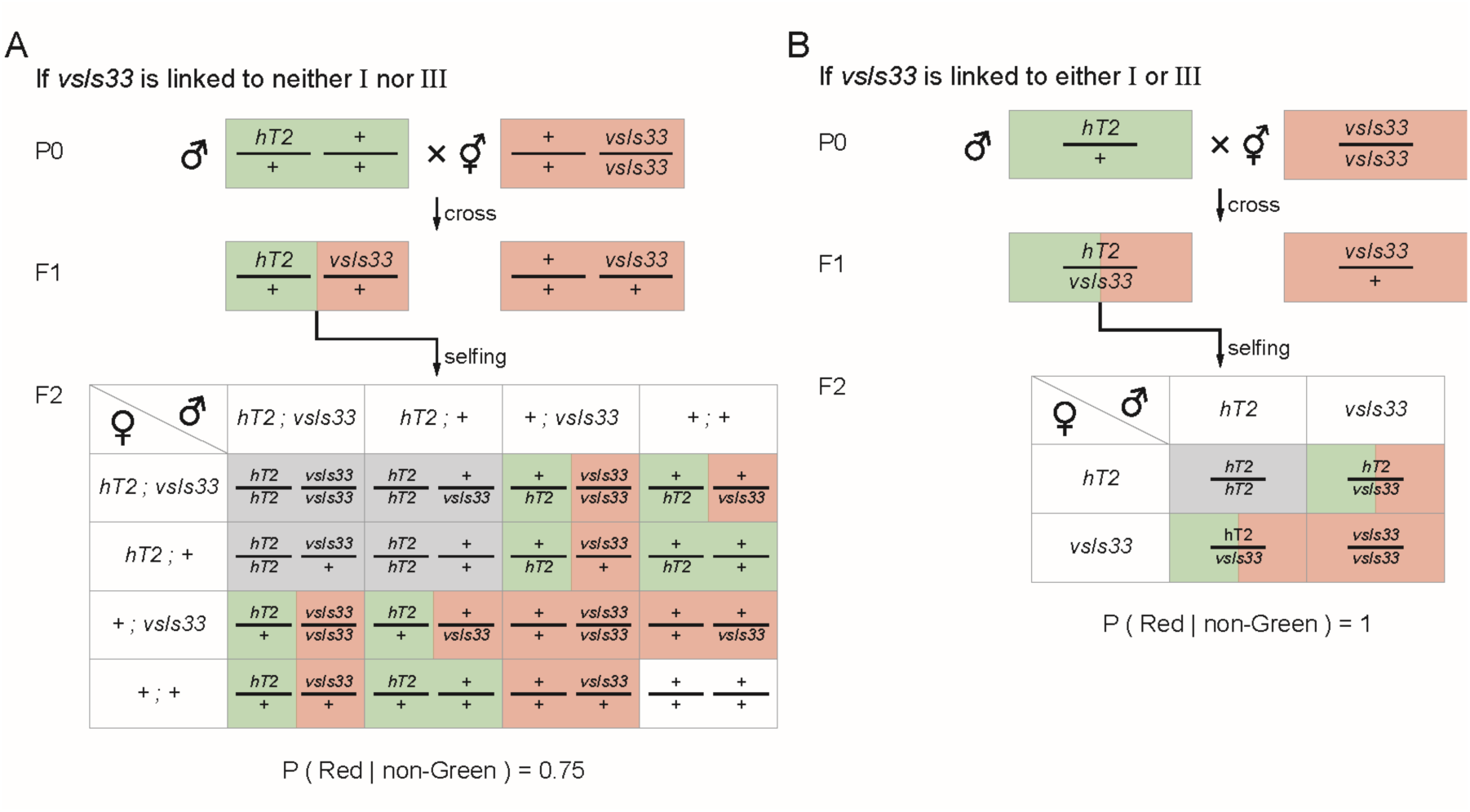
Theory of the I/III-linkage test. *(A)* The Punnett square showing the F2s to be observed if *vsIs33* is linked to neither I nor III. *hT2* is homozygous lethal. *(B)* The Punnett square showing the F2s to be observed if *vsIs33* is linked to either I or III. Green: green fluorescent animal. Red: red fluorescent animal. Gray: inviable animal.

**Fig. S9.**
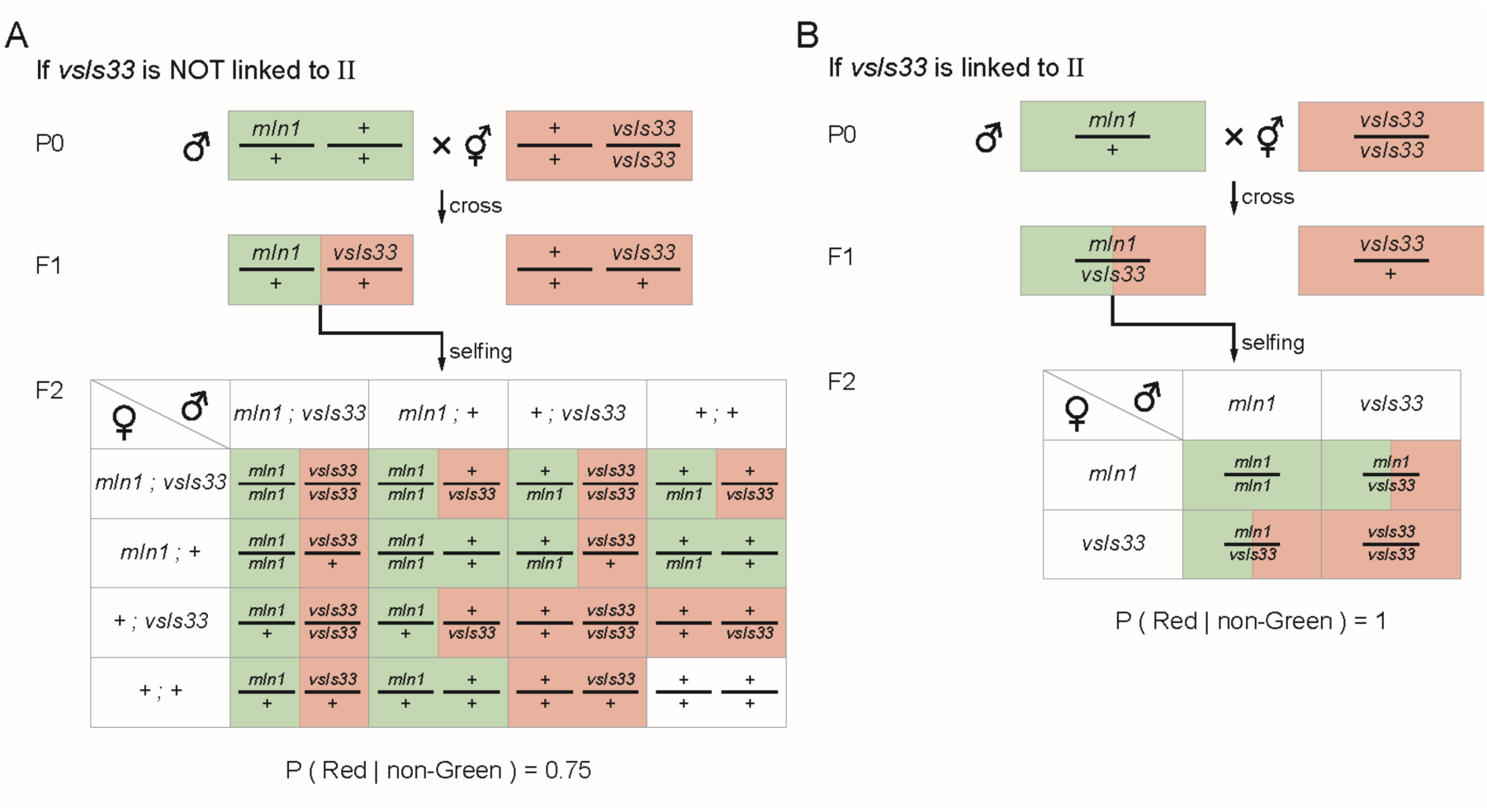
Theory of the II-linkage test. *(A)* The Punnett square showing the F2s to be observed if *vsIs33* is not linked to II. *(B)* The Punnett square showing the F2s to be observed if *vsIs33* is linked to II. The color code is the same as in Fig. S8.

**Fig. S10.**
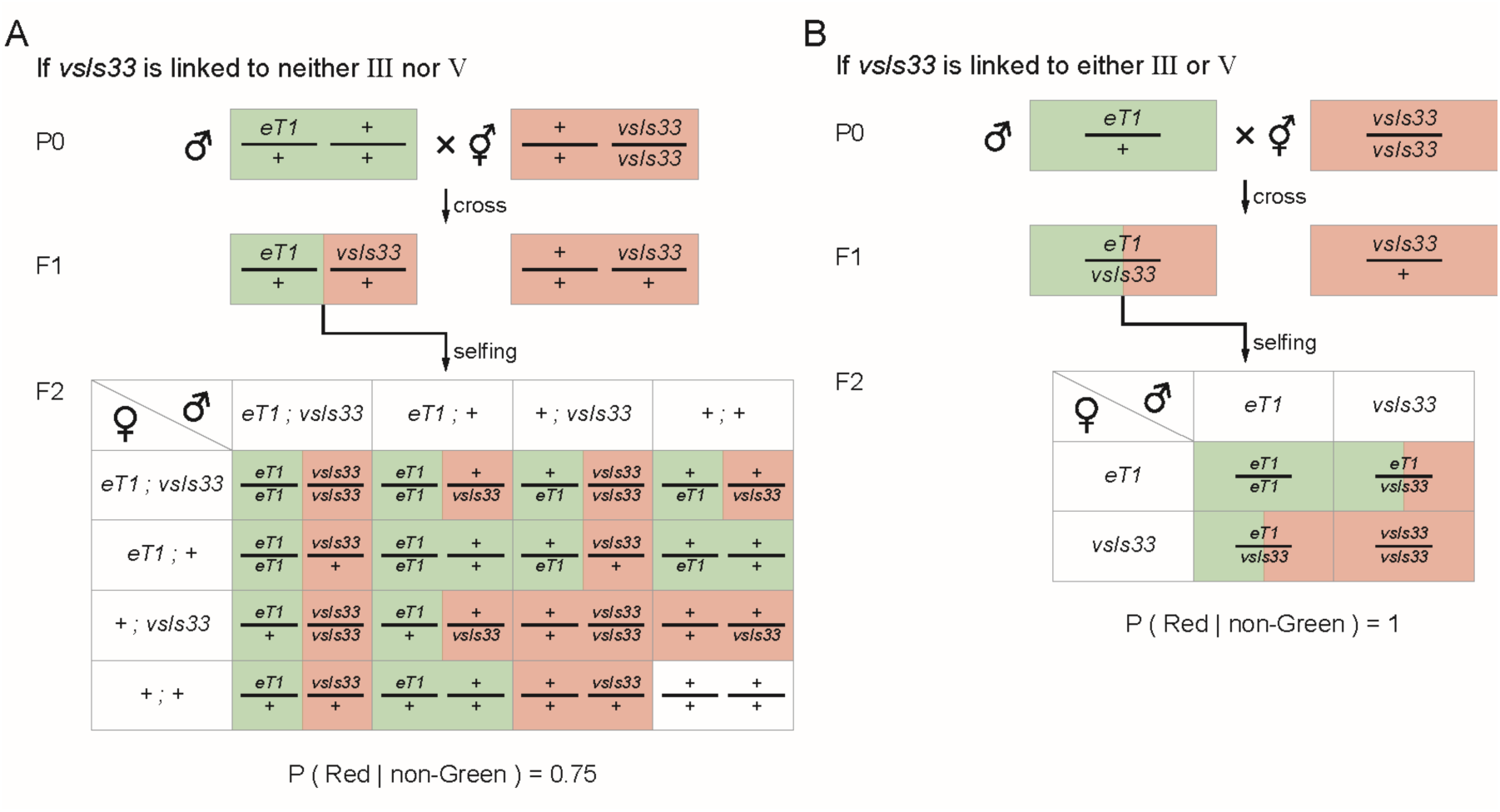
Theory of the III/V-linkage test. *(A)* The Punnett square showing the F2s to be observed if *vsIs33* is linked to neither III nor V. *(B)* The Punnett square showing the F2s to be observed if *vsIs33* is linked to either III or V. The color code is the same as in Fig. S8.

**Fig. S11.**
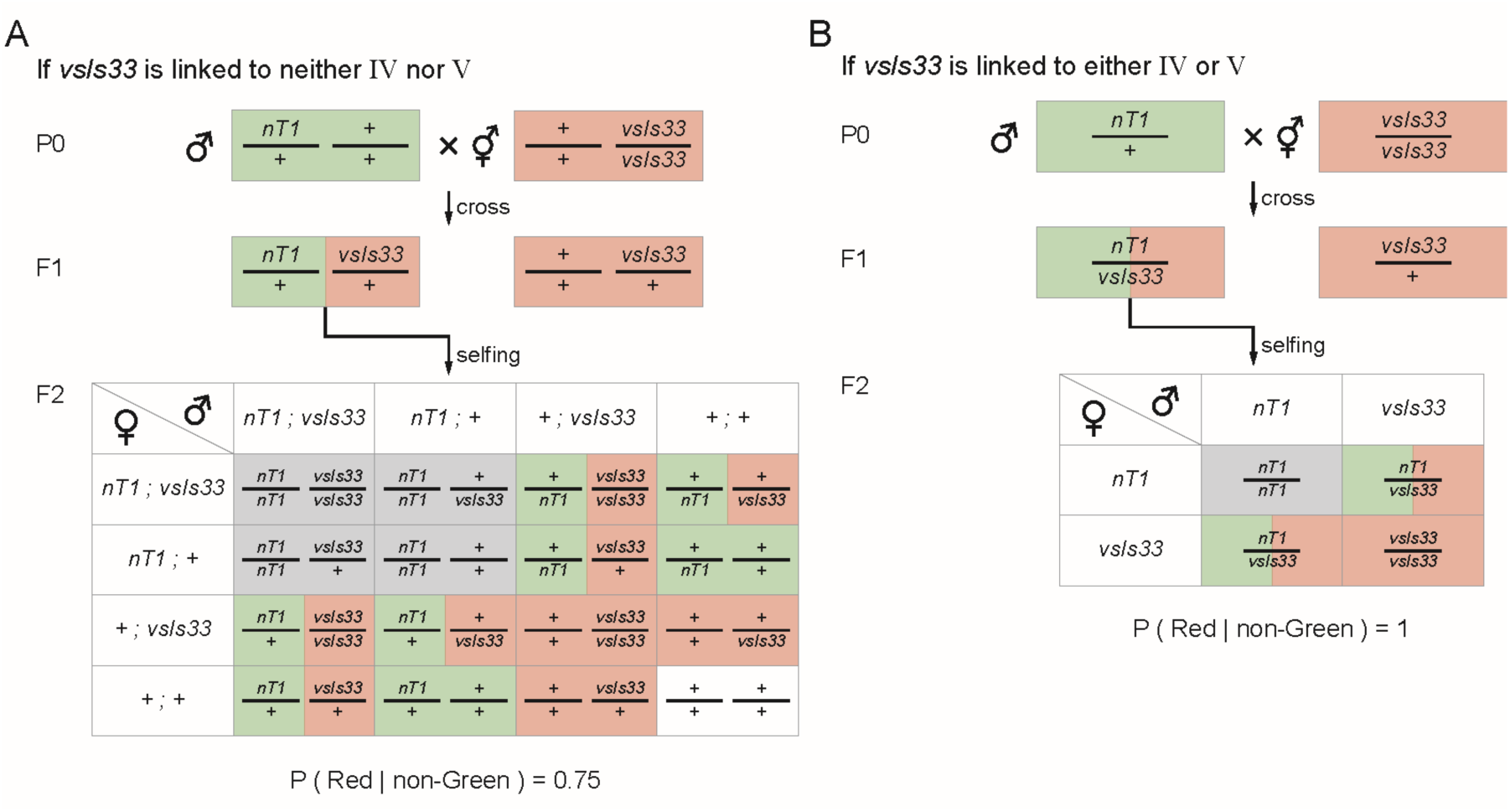
Theory of the IV/V-linkage test. *(A)* The Punnett square showing the F2s to be observed if *vsIs33* is linked to neither IV nor V. *nT1* is homozygous inviable. *(B)* The Punnett square showing the F2s to be observed if *vsIs33* is linked to either III or V. The color code is the same as in Fig. S8.

**Fig. S12.**
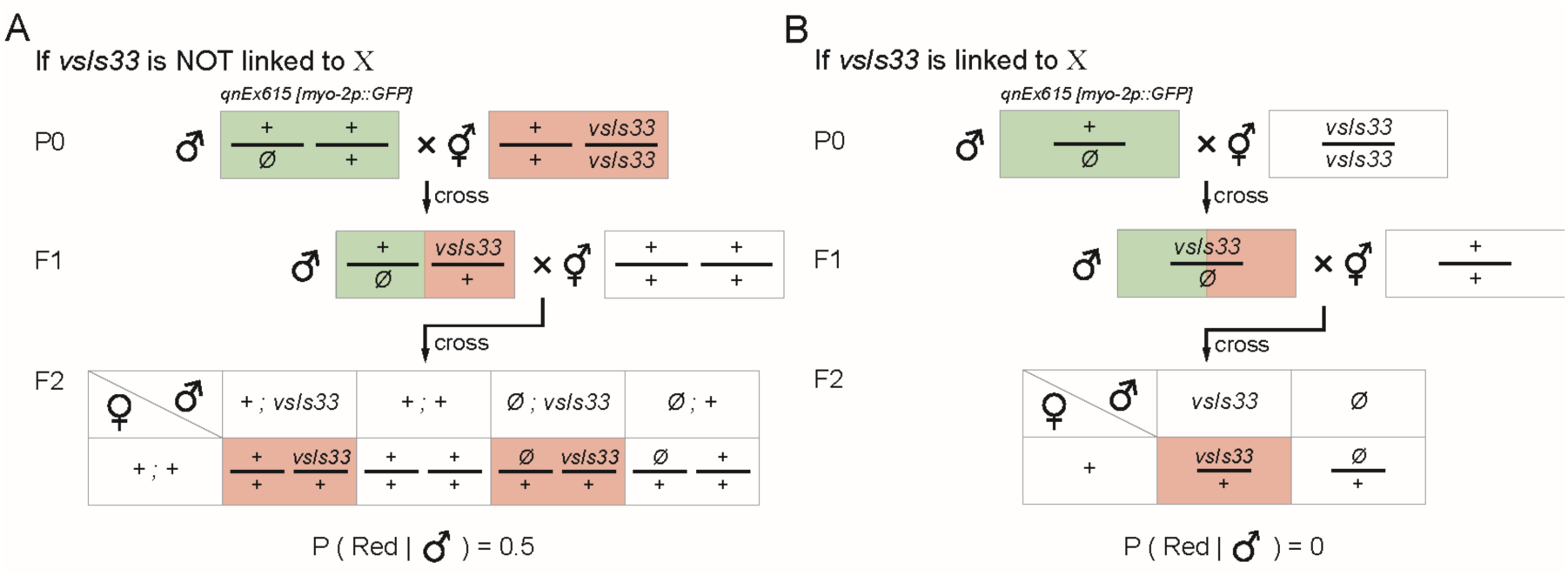
Theory of the X-linkage test. *(A)* The Punnett square showing the F2s to be observed if *vsIs33* is not linked to X. *(B)* The Punnett square showing the F2s to be observed if *vsIs33* is linked to X. The color code is the same as in Fig. S8.

**Fig. S13.**
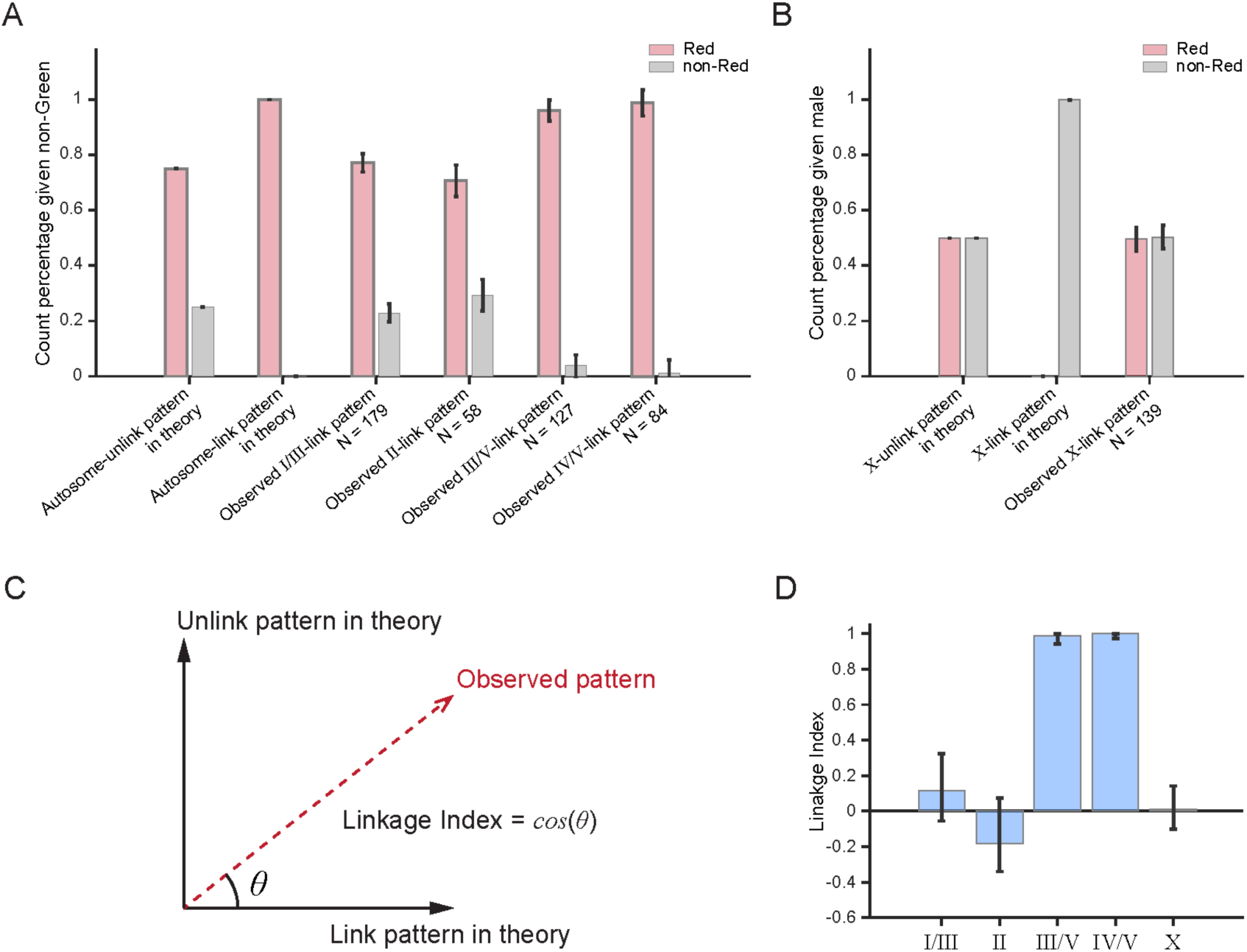
Data of the linkage test for *vsIs33*. *(A)* The theory predicted and the observed count percentages of Red and non-Red among the non-Green F2s for the autosome-linkage test. Error bar: standard deviation of the observed percentages under the autosome-unlink assumption. *(B)* The theory predicted and the observed count percentages of Red and non-Red among the F2 males for the X-linkage test. Error bar: standard deviation of the observed percentages under the X-unlink assumption. *(C)* Graphical representation of Linkage Index. *(D)* Observed Linkage Indices for *vsIs33* over different chromosomes. Error bar: standard deviation of the observed Linkage Indices under the unlink assumption.

**Table S1.** Viability data for the animals 24 hours after the WormPicker and manual picking. ***p***_***a***_, ***p***_***d***_, and ***p***_*e*_ denote the percentage of the animal alive, dead, and escaped; ***N*** denotes the number of animals picked.

| Genotype | Phenotype | Stage | Automated picking |  |  |  | Manual picking |  |  |  |
| --- | --- | --- | --- | --- | --- | --- | --- | --- | --- | --- |
| | | | $P_a$ | $P_d$ | $P_e$ | $N$ | $P_a$ | $P_d$ | $P_e$ | $N$ |
| N2 | Wild type | L1 | 90.7 | - | - | 54 | 98 | 0 | 2 | 50 |
|  |  | L2 | 92.8 | - | - | 83 | 98 | 0 | 2 | 50 |
|  |  | L3 | 96.7 | 0 | 3.3 | 90 | 98 | 0 | 2 | 50 |
|  |  | L4 | 98.6 | 0 | 1.4 | 145 | 100 | 0 | 0 | 50 |
|  |  | Day 1 adult | 94.3 | 0 | 5.7 | 105 | 100 | 0 | 0 | 60 |
|  |  | Day 5 adult | 96.4 | 3.6 | 0 | 84 | 97.1 | 2.9 | 0 | 35 |
| *lin-15* | Multivulva | Adult | 81.5 | 17 | 1.5 | 65 | 92 | 8 | 0 | 50 |
| *rol-6* | Roller | L4 - Adult | 98 | 0 | 2 | 50 | 100 | 0 | 0 | 61 |
| *unc-13* | Uncoordinated, paralyzed | L3 - Adult | 100 | 0 | 0 | 53 | 100 | 0 | 0 | 50 |
| *lpr-1* | Fragile cuticles | L4 - Adult | 98.1 | 0 | 1.9 | 54 | 98 | 2 | 0 | 50 |

**Table S2.**
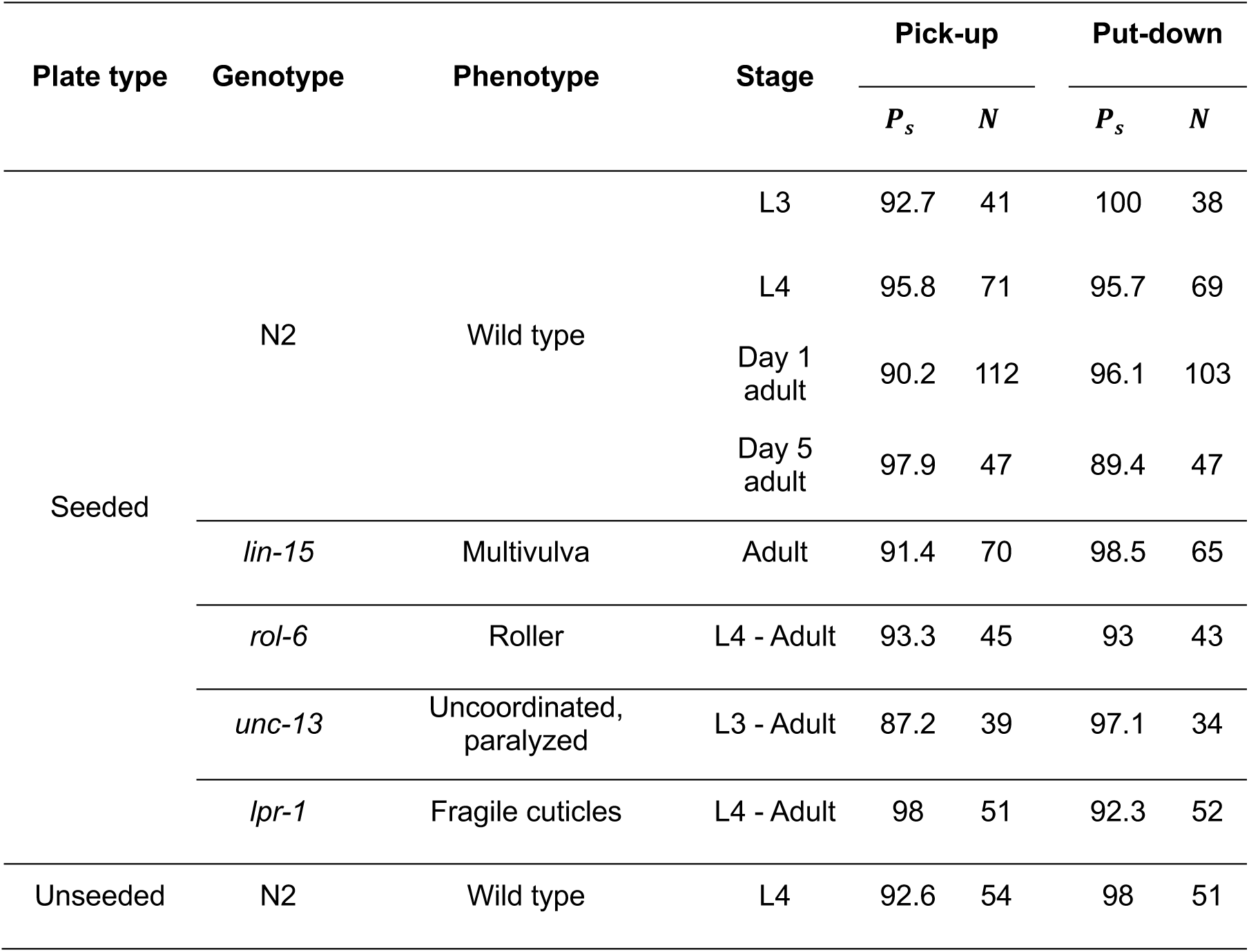
Success rates for WormPicker picking up and putting down *C. elegans*. ***p***_***s***_ denotes the percentage of the successful attempts; ***N*** denotes the total number of the attempts made.

**Table S3.**
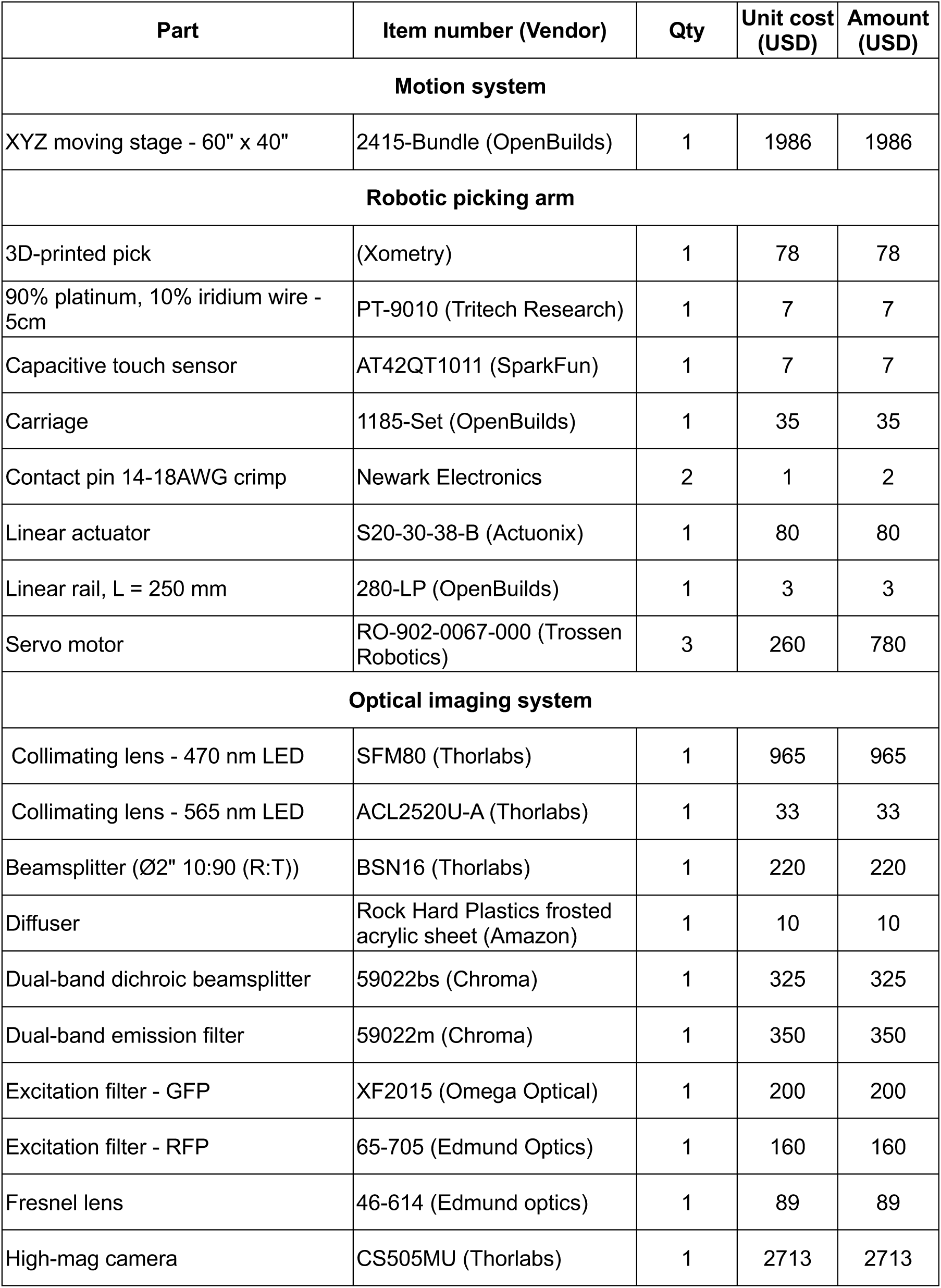
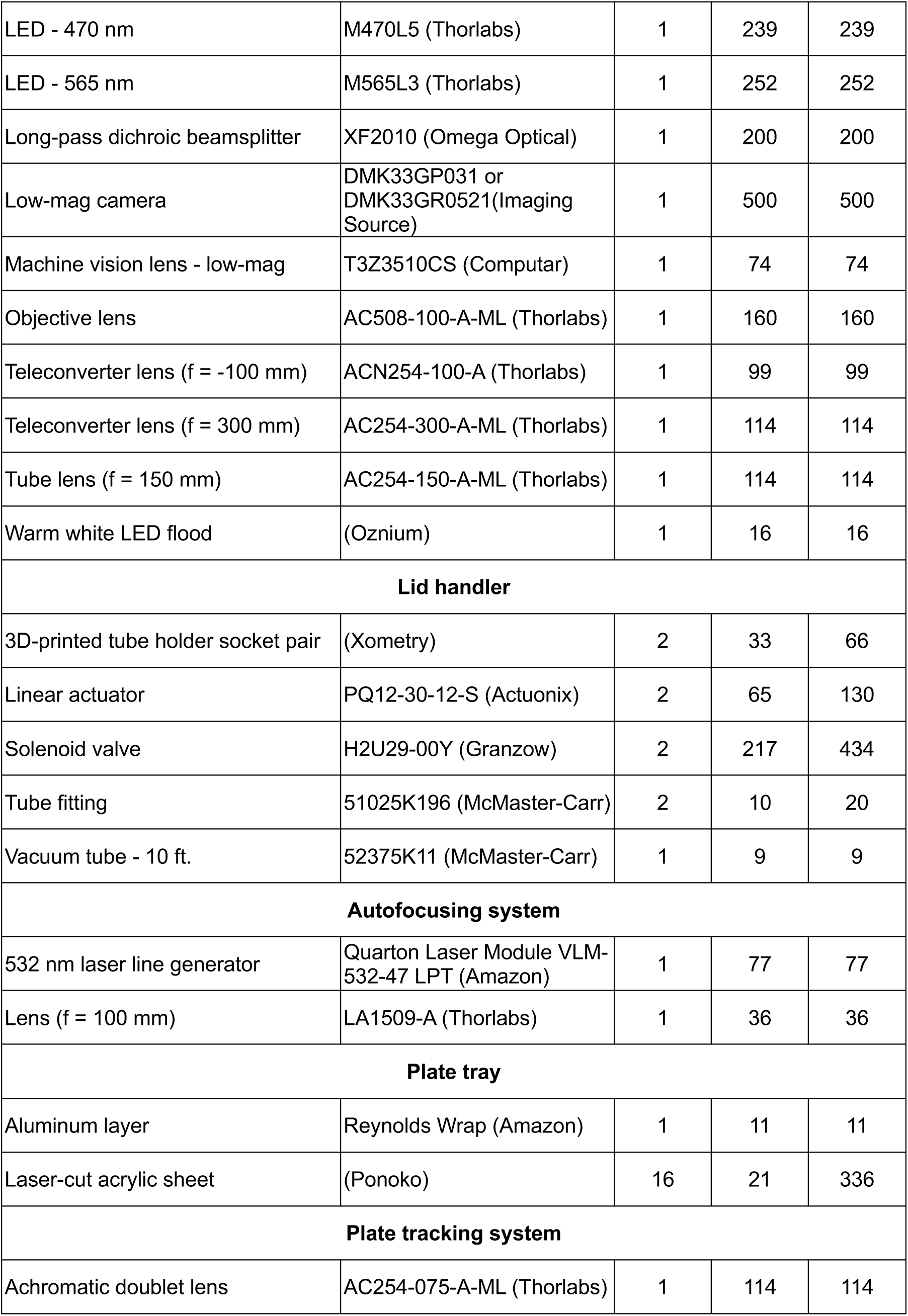
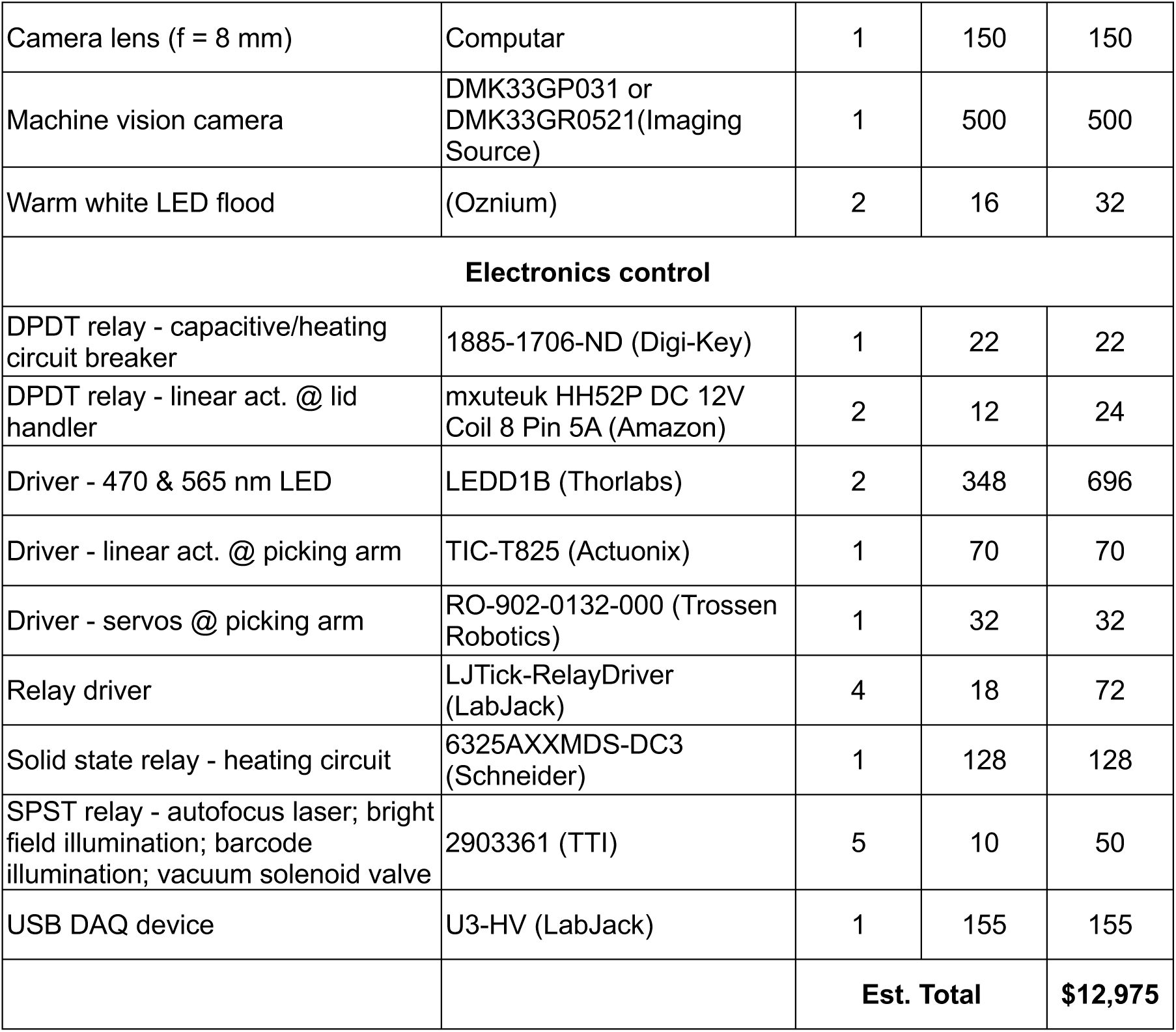
WormPicker key components list

**Table S4.** Fluorescence imaging and processing parameters. ***T***_*e*_, ***h***_***i***_, and ***h***_***s***_ denote the exposure time, intensity threshold, and size threshold.

| Experiment | Purpose | Fluorescent transgene of interest | Imaging and processing parameters |  |  |
| --- | --- | --- | --- | --- | --- |
| | | | $T_e$<br>(ms) | $h_i$<br>(a. u.) | $h_s$<br>(pixels) |
| Genetic cross<br>(Fig. 3) | Assay transgene expression | *vsIs28 [dop-1p::GFP]* | 200 | 70 | 20 |
| Genetic mapping<br>(Fig. 4) | Test I/III-linkage | *vsIs33 [dop-3p::RFP]* | 100 | 80 | 20 |
|  |  | *hT2 {qls48 [myo-2::GFP]}* | 200 | 80 | 20 |
|  | Test II-linkage | *vsIs33 [dop-3p::RFP]* | 100 | 80 | 50 |
|  |  | *mIn1 {mls14 [myo-2::GFP]}* | 200 | 60 | 50 |
|  | Test III/V-linkage | *vsIs33 [dop-3p::RFP]* | 100 | 60 | 20 |
|  |  | *eT1 {umnIs12 [myo-2::GFP]}* | 200 | 130 | 20 |
|  | Test IV/V-linkage | *vsIs33 [dop-3p::RFP]* | 100 | 80 | 20 |
|  |  | *nT1 {qls51 [myo-2::GFP]}* | 200 | 80 | 20 |
|  | Test X-linkage | *vsIs33 [dop-3p::RFP]* | 100 | 80 | 20 |
|  |  | *qnEx615 [myo-2p::GFP]* | 200 | 80 | 20 |
| Genomic integration<br>(Fig. 5) | Assay transgene expression | *qhEx265 [unc-47p::GFP]* | 200 | 70 | 1 |

**Table S5.** Pick motion trajectory parameters. *v*_***z***_ and ***D***_***z***_ denote the velocity and the displacement of the component in ***z*** direction; ***α*** and ***δ*** denote the angular velocity and the angular displacement of the servo motor.

| Phase | Linear actuator |  | Servo 1 |  | Servo 2 |  | Servo 3 |  | Main gantry |  |
| --- | --- | --- | --- | --- | --- | --- | --- | --- | --- | --- |
| | $v_z$<br>(mm/s) | $D_z$<br>(mm) | $\alpha$<br>(deg/s) | $\delta$<br>(deg) | $\alpha$<br>(deg/s) | $\delta$<br>(deg) | $\alpha$<br>(deg/s) | $\delta$<br>(deg) | $v_z$<br>(mm/s) | $D_z$<br>(mm) |
| **Picking up** |  |  |  |  |  |  |  |  |  |  |
| i | -20 | -2.5 | 0 | 0 | 0 | 0 | 0 | 0 | 0 | 0 |
| ii | 0 | 0 | 0 | 0 | 274.8 | 1.23 | 0 | 0 | 0 | 0 |
| iii | 0 | 0 | 274.8 | 0.44 | 0 | 0 | 274.8 | 0.44 | 0 | 0 |
| **Putting down** |  |  |  |  |  |  |  |  |  |  |
| i | -20 | -2.5 | 0 | 0 | 0 | 0 | 0 | 0 | 0 | 0 |
| i-ii<br>waiting<br>(s) | 1 |  |  |  |  |  |  |  |  |  |
| ii | 0 | 0 | 0 | 0 | -274.8 | -1.58 | 0 | 0 | 0 | 0 |
| ii-iii<br>waiting<br>(s) | 2 |  |  |  |  |  |  |  |  |  |
| iii | 0 | 0 | 0 | 0 | 0 | 0 | 0 | 0 | -125 | -0.4 |
| iv | 0 | 0 | 274.8 | 39.11 | 0 | 0 | 0 | 0 | 0 | 0 |

**Table S6.** Descriptions of the source scripts in WormPickerLib.

| Category | Name | Description |
| --- | --- | --- |
| High-level | CrossWorms | Pick hermaphrodites and males matching the specified phenotypes from two source plates to one destination plate. |
|  | ScreenPlates | Screen multiple plates for assaying population phenotypes. |
|  | SingleWorms | Pick worms matching the specified phenotypes to individual plates. A single worm per plate. |
| Mid-level | PickNWorms | Pick multiple worms matching the specific phenotypes from one plate to another. |
|  | ScreenOnePlate | Screen a single plate for assaying population phenotypes. |
| Low-level | AutoFocus | Focus the image. |
|  | CalibOx | Calibrate how many pixel-translations in the image corresponding to one unit movement in the gantry. |
|  | CalibPick | Calibrate the pick position in the low-magnification FOV. |
|  | CalibTray | Calibrate the plate positions over the plate tray platform. |
|  | CenterWorm | Track worms and move individual animals to the high-magnification FOV. |
|  | HeatPick | Sterilize the pick. |
|  | MoveToPlate | Move the gantry to right above a plate. |
|  | Phenotype | Phenotype any worms found in the high-magnification FOV. |
|  | ReadBarCode | Read the barcode identifier for a plate, write/read the plate information to/from the database. |
|  | SpeedPick | Perform a single pick-up/put-down action. |

**Table S7.**
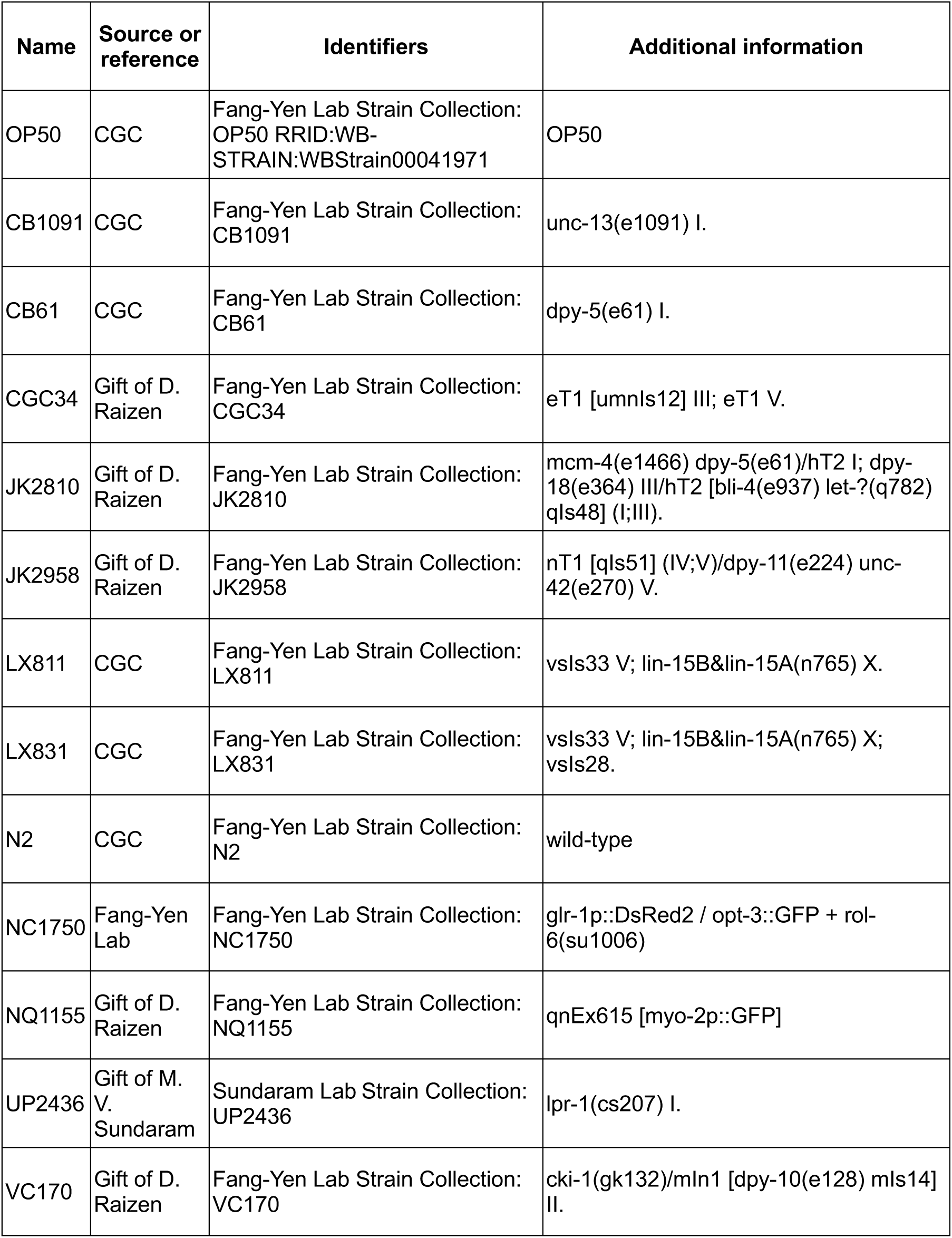
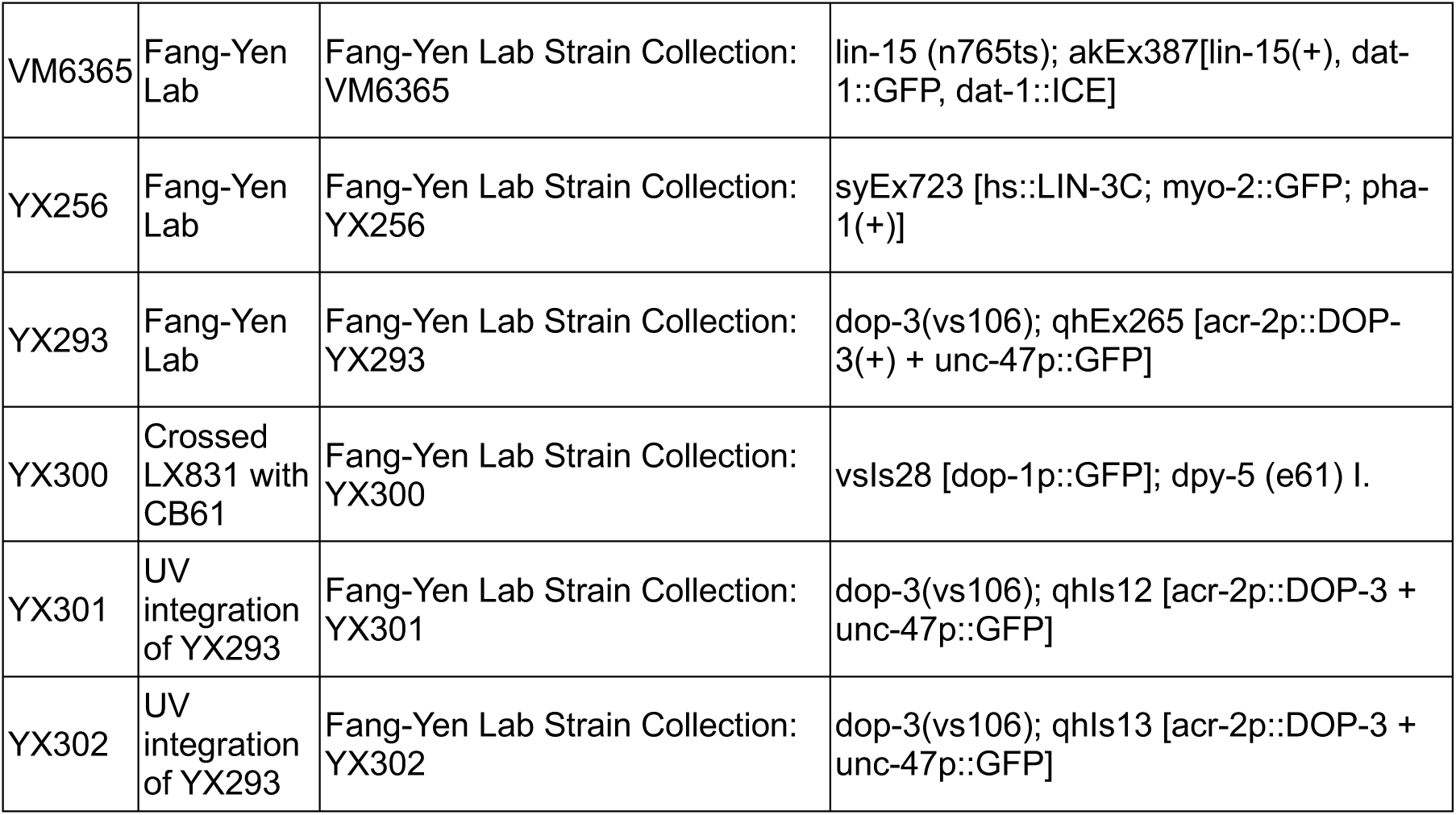
Strains used in this study.

**Movie S1 (separate file).** Automated multimodal imaging of *C. elegans.* Transgenic (JK2958) *C. elegans* in bright field images followed by an image from the GFP channel. Transgenic (LX811) *C. elegans* in bright field images followed by an image from the RFP channel. FOV: 1.88 mm x 1.57 mm.

**Movie S2 (separate file).** Lid manipulation. Lids of agar plates on the platform are removed and replaced using two motorized vacuum actuators.

**Movie S3 (separate file).** Pick sterilization. The platinum wire loop pick is sterilized by resistive heating.

**Movie S4 (separate file).** Automated *C. elegans* picking from the WormPicker’s camera view. Overlay of the high-magnification and low-magnification video streams. A worm is automatically picked up by the wire loop pick and transferred to a different plate.

**Movie S5 (separate file).** Automated *C. elegans* transfer between agar substrates. An electric current sterilizes the pick; worms are tracked and phenotyped; the robotic arm picks up a worm from a plate and transfers it to a second plate.

**Dataset S1 (separate file).** Source data. The file contains the source data for the plots presented in this paper.

**Design File S1 (separate file).** WormPicker mechanical design file. The file contains a CAD design (F3D format) for the WormPicker hardware system, including a plate tray platform, X, Y, Z linear carriage assemblies, an optical imaging system, two lid handlers, a robotic picking arm, an autofocusing system, an illuminator (under the platform), and a plate tracking system (under the platform).

